# Lifespan trajectory of chimpanzee brains characterized by magnetic resonance imaging histology

**DOI:** 10.1101/2024.12.06.627145

**Authors:** I. Lipp, E. Kirilina, C. Jäger, M. Morawski, A. Jauch, K.J. Pine, L.J. Edwards, S. Helbling, D. Rose, G. Helms, C. Eichner, T. Deschner, T. Gräßle, EBC Consortium, P. Gunz, A. Anwander, A.D. Friederici, R.M. Wittig, C. Crockford, N. Weiskopf

**Affiliations:** Department of Neurophysics, Max Planck Institute for Human Cognitive and Brain Sciences, Leipzig, Germany; Center for Computational Neuroscience, Free University Berlin, Berlin, Germany; Paul Flechsig Institute - Center of Neuropathology and Brain Research, Faculty of Medicine, Universität Leipzig, Germany; Ernst Strüngmann Institute for Neuroscience in Cooperation with Max Planck Society, Frankfurt am Main, Germany; Department of Clinical Sciences and Medical Radiation Physics, Lunds Universitet, Lund, Sweden; Department of Neuropsychology, Max Planck Institute for Human Cognitive and Brain Sciences, Leipzig, Germany; Institute for Cognitive Sciences, University of Osnabrueck, Osnabrueck, Germany; Ozouga Chimpanzee Project, Loango NP, Gabon; Epidemiology of Highly Pathogenic Microorganisms, Robert Koch Institute, Berlin, Germany; Department of Human Evolution, Max Planck Institute for Evolutionary Anthropology, Leipzig, Germany; Department of Human Behavior, Ecology and Culture, Max Planck Institute for Evolutionary Anthropology, Leipzig, Germany; Ape Social Mind Lab, Institute of Cognitive Science Marc Jeannerod, UMR 5229, CNRS, Lyon, France; Taï Chimpanzee Project, Centre Suisse de Recherches Scientifiques, Abidjan, Cote d’Ivoire; Felix Bloch Institute for Solid State Physics, Faculty of Physics and Earth Sciences, Leipzig University, Leipzig, Germany

## Abstract

Chimpanzee brain maturation provides an invaluable framework for understanding the evolution of the human brain. We performed ultra-high resolution quantitative magnetic resonance imaging (qMRI) with histological validation on *post mortem* brains from captive and wild chimpanzees with a broad age range. We mapped developmental myelination and age-related iron accumulation across regions and layers of the neocortex. Compared to humans, chimpanzees showed more myelination and iron deposition in motor and premotor cortices, while the auditory cortex was more strongly myelinated in humans. Our model suggests that chimpanzees’ cortical myelination was largely completed by the age of nine years, while iron accumulation continued throughout the lifespan. The regions with highest adult levels of myelin and iron took the longest to mature, challenging the widespread assumption that highly myelinated regions complete their development first. The reported maps and developmental curves provide a foundation for comparative neuroscience research and understanding of human brain evolution.

## Introduction

The evolution of unique human skills, such as syntax-based language^1^, sophisticated tool use^2^, and complex social cognition^3^ came with the expansion and differentiation of the hominin brain. Fossil records show that humans evolved from australopiths with brains that had chimpanzee-like folding patterns, and adult endocranial volumes that were on average 20% higher than in chimpanzees^4^. The brains of chimpanzees differ from human brains substantially on the macroscopic and microscopic levels with differences in size and cortical surface area^5^, prefrontal cortex morphology^6^, neocortical dendritic morphology^7^ and projections of white matter tracts supporting language processing in humans^8^. These differences in neuroanatomy may be the result of divergent ontogenetic trajectories between species. Cross-species comparison of brain lifespan trajectories on the macro- and microscale^5,9–11^ is indispensable to identify the evolutionary changes that facilitated the development of skills in the more recent stages of hominin evolution or contributed to the emergence of human-specific pathologies associated with certain developmental periods^12^. However, our knowledge about macroscopic and microscopic lifespan trajectories of great ape brains are limited, and data which can be compared between human and great apes are scarce, mostly due to methodological and ethical limitations.

Previous ontogenetic studies in chimpanzees using Magnetic Resonance Imaging (MRI) have focused on macroscopic measures of gray and white matter volumes. They suggest less rapid early development of the prefrontal white matter in chimpanzees^11^, less aging-related changes in neocortical gray and white matter in chimpanzees^9^ and negligible developmental changes in gray and white matter volume between ages 8 and 53^10^. However, volume changes are macroscopic measures, which reflect and conflate many factors^13^, as brain ontogeny involves many processes that shape the brain, on both the macroscopic and microscopic level^14,15^.

Here, we characterized the microscopic changes in the chimpanzee neocortex using *post mortem* quantitative MRI at ultra-high resolutions by using ethically and sustainably sourced brains from chimpanzees after their death in captivity or in the wild. We quantified two major processes in the brains across the lifespan - developmental myelination and age related iron accumulation.

Myelination is a developmental process that starts before birth^16,17^ and continues into adulthood^18^. In the cortex, the level of myelin can provide information on the functional specialization of cortical areas. Primary sensory and motor areas are myelinated more highly and presumably earlier in development than association areas^19–21^. Low cortical myelination seems to be a factor beneficial to cortical plasticity^20^, and areas low in myelin expanded most during evolution from macaques to humans, as well as during ontogenetic development^22^ and are particularly sensitive to neurodevelopmental abnormalities^23^. Not only the total cortical myelin content changes with development, but also myelination patterns across cortical layers^24,25^.

One postulated characteristic feature of human brain development is a prolonged period of cortical myelination compared to chimpanzees^26^, particularly pronounced in higher level areas such as the prefrontal cortex. However, previous studies on cortical myelination in chimpanzees are limited to manually analyzed histological tissue sections with stereology from only a small subset of cortical regions^26^ and have not considered layer-dependency of cortical myelination.

Another hallmark of brain lifespan trajectory is iron accumulation with age. In the brain, iron plays a dual role. On the one hand, iron is a co-factor facilitating crucial processes in the brain including myelin synthesis, energy production and neurotransmitter synthesis^27,28^. Due to its involvement in myelination, iron often colocalizes with myelin in the cortex^29^. On the other hand, when present in high concentrations, highly reactive iron is a source of oxidative stress and mitochondrial damage^30^. Iron overload in age is an important contributor to neurodegeneration, however, a sufficient level of iron is indispensable for healthy brain development and function^29^. Therefore, comparison of life span iron accumulation between species is a key step towards understanding differences in brain development and aging.

MRI has become the major neuroimaging method for mapping brain anatomy and development in humans^31^ and a cornerstone of comparative neuroscience^32^. Recent advances in quantitative MRI have enabled the characterization of brain microstructure^33,34^ and developmental trajectories^25,35,36–38^. Multi-parametric MRI mapping is particularly powerful when studying biologically complex processes such as brain development^35,39,40^, since it yields multiple parameters with differing microstructural sensitivities. When used at high spatial resolutions, quantitative MRI can depict the layer-dependent levels of myelin and iron in the cortex^24,25,34^.

To date, the existing MRI data of chimpanzee brains have been limited to non-quantitative scans performed *in vivo* on anesthetized animals in primate research centers^6,41^. Due to ethical concerns, application of these invasive methods to great apes has largely been halted. The existing datasets are unlikely to be extended and cannot profit from recent developments in MRI methodology. Here, we showcase how to conduct neuroscientific research in primates with *post mortem* quantitative MRI at ultra-high resolutions by using ethically and sustainably sourced brains from chimpanzees after their deaths in captivity or the wild^42^.

We provide whole-brain MRI-based cortical myelination and iron reference maps from a group of chimpanzees. Specific spatio-temporal ontogenetic changes were observed in the myelin and iron markers as well as in intracortical profiles. In addition to the new insights into ontogenetic development, our work shows that *post mortem* studies could bring a step change to the field of comparative neuroscience that has so far been hindered by the practical and ethical implications of obtaining *in vivo* data.

## Results

We have established an ethically sustainable pipeline to collect *post mortem* brains of wild and captive chimpanzees^42^, acquire whole brain 300 μm resolution quantitative MRI data, process them with optimized data analysis strategies^43^ and conduct histological and immunohistochemical staining and microscopic imaging (**Figure 1**). Altogether, 20 chimpanzee brains with ages spanning the entire chimpanzee lifespan were obtained from the network of African field sites, zoos, and sanctuaries in the period from 02.2018 to 08.2020. Data from 17 brains were analyzed (10 wild, 6 captive, 1 sanctuary; 11 male, 6 female; age [mean *±* sd]: 23 *±* 19 years, details for all brains are shown in Supplementary Table 1). Three brains were excluded from all analyses due to heterolytic damage of brain tissue induced by a systemic bacterial infection, which was the cause of death in these three animals. Ages at death (± uncertainty for ages that had to be estimated^44,45^) of the 17 animals used in the analysis were: 0.1, 1 (±1), 1.6, 1.75, 2.75, 6, 12 (±3), 13 (±3), 17.5 (±5), 30 (±5), 34, 40 (±5), 43 (±5), 44, 45 (±5), 47, 52 years.

**Figure 1:**
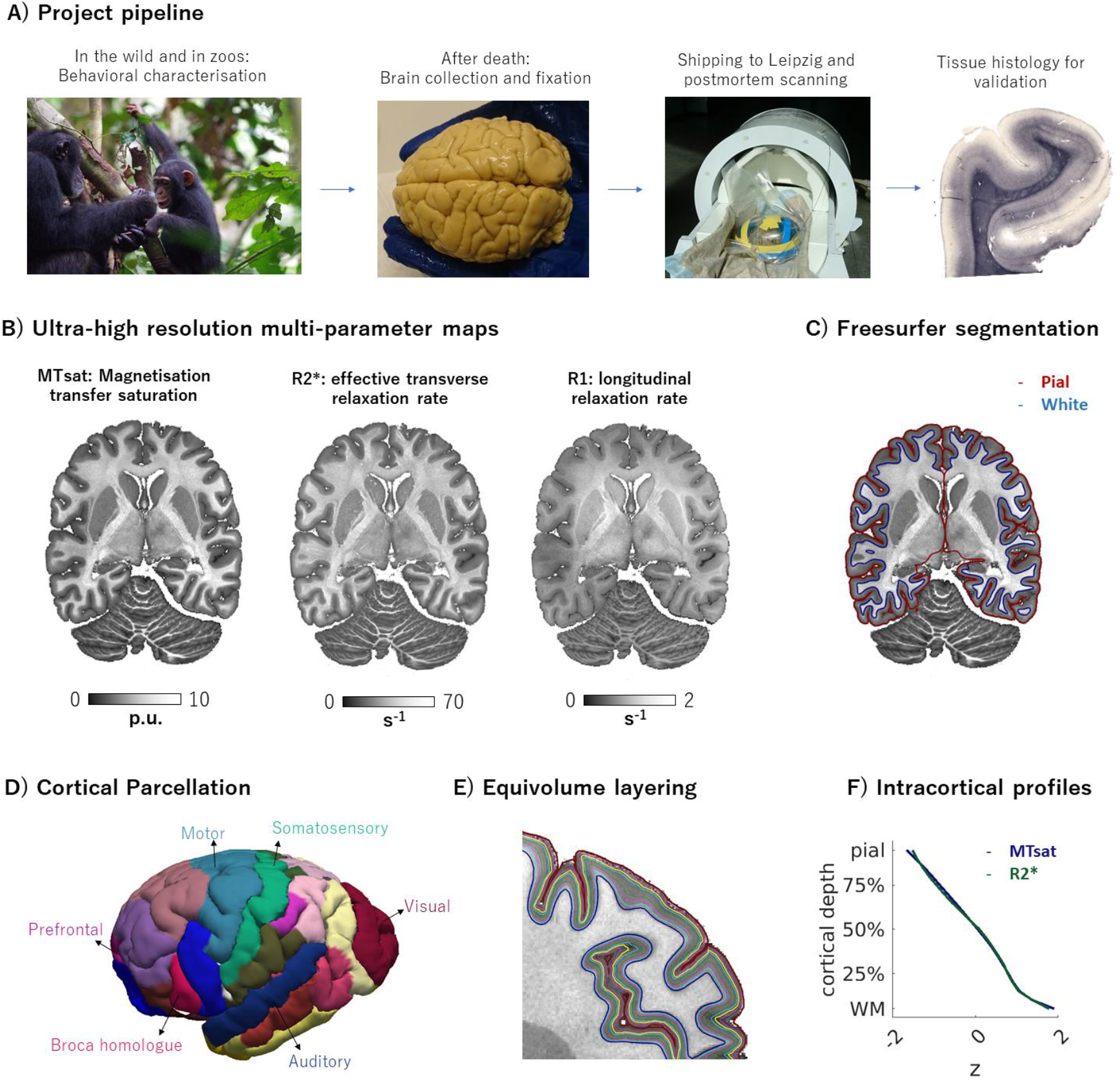
**Overview of the study workflow and main outcomes**. **A.** Chimpanzees that live in the wild or in zoos are behaviourally characterized and observed by researchers (data not reported here). After death, veterinarians/pathologists extract the brain and fix it in formalin before shipping it to Leipzig, Germany, for *post mortem* MRI and subsequent histology for microscopical validation. **B.** Ultra-high resolution quantitative parameter mapping is performed at 7T. Example quantitative parameter maps of one chimpanzee are shown. **C.** Freesurfer is used to obtain pial and white matter surfaces of individual brains. **D.** The cortex is parcellated using the BB38 chimpanzee atlas, and regions of interest are defined based on this parcellation (the colors used here to visually separate the regions are arbitrary). **E.** An equivolume-layering approach is used to sample the quantitative parameters from different cortical depths and (**F**) to produce intracortical profiles of quantitative MRI parameters.

### Ultra-high resolution qMRI metrics of myelin and iron in chimpanzee brains

Whole-brain maps of quantitative MRI parameters obtained on a 7T human scanner at an ultra-high isotropic resolution of 300 μm revealed exquisite anatomical details within the cortex, white matter and subcortex, clearly demonstrating layer-dependent intracortical contrast. QMRI provided three quantitative whole brain metrics which reflect tissue myelination and iron-content (**Figure 2** **top**), and can be compared across cortical layers, across brain areas, across individuals and across species. The magnetization transfer saturation MTsat is a sensitive and specific marker for cortical myelination^46^ while the effective transverse relaxation rate (R2*) and longitudinal relaxation rate (R1) are sensitive to myelin and iron content^47,48^. High quality R2* and MTsat maps were obtained for the 17 brains. R1 maps in 3 of 17 brains were excluded from analyses as they showed some artifacts probably related to autolytic damage.

**Figure 2:**
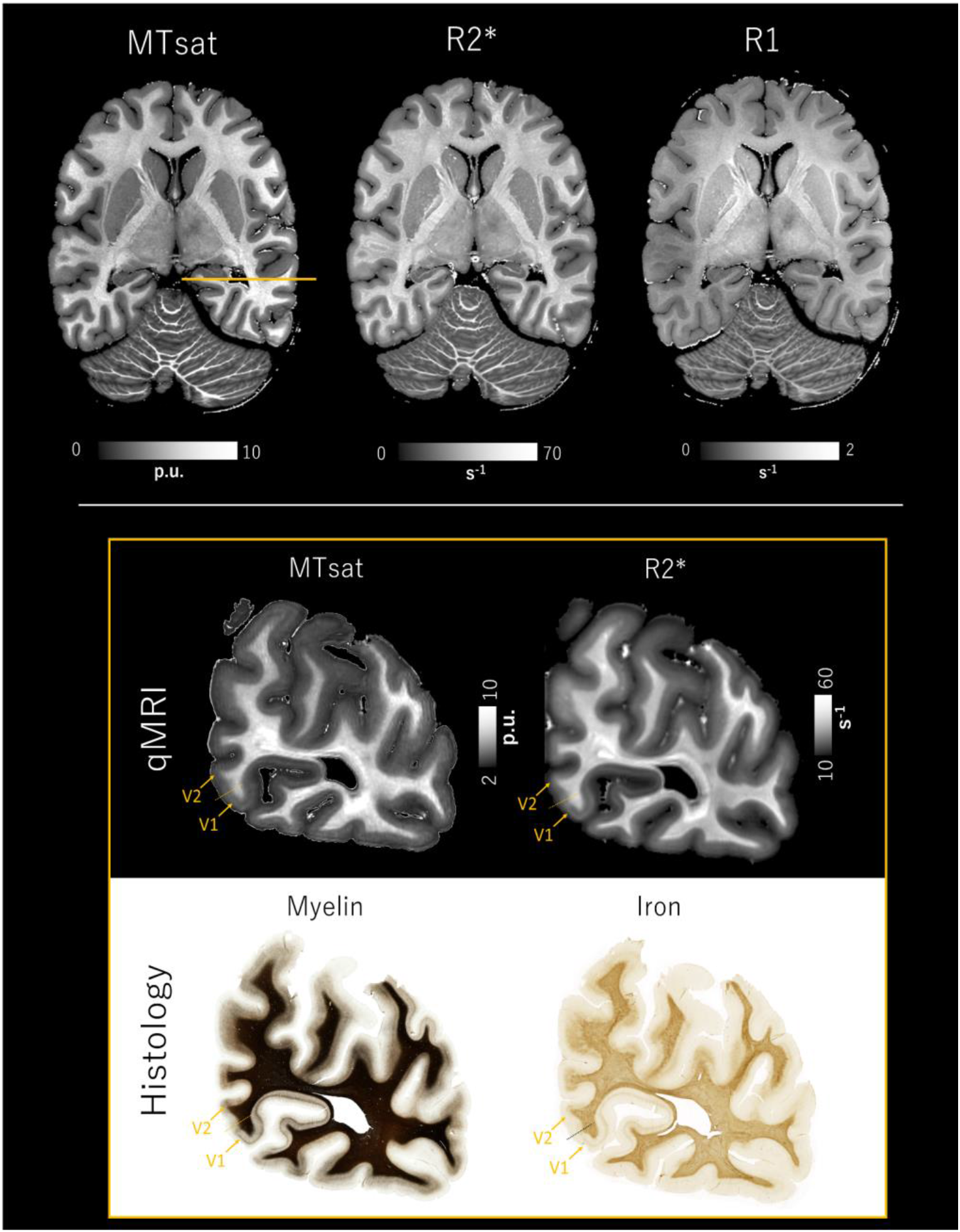
Top: Example 300 μm high resolution quantitative MRI data set from a wild, 45 year old male chimpanzee’s brain. Shown is an axial slice for each of the three quantitative contrasts: Magnetization transfer saturation (MTsat), effective transverse relaxation rate (R2*), and longitudinal relaxation rate (R1). All parameters show an exquisite contrast between gray and white matter. The high resolution allows identification of detailed neuroanatomical features (e.g., cortical sheet, basal ganglia, cerebellar folia) and the assessment of differences in microstructure between cortical areas and across cortical depth. Note that the R1 map appears brighter in the center of the brain which is likely an artifact of fixation. **Bottom:** Quantitative MRI parameters MTsat (myelin-sensitive) and R2* (myelin- and iron-sensitive) validated by histological stainings of coronal sections of the occipital lobe from a sanctuary-housed, 12 year old male chimpanzee. Myelin was visualized with a Gallyas silver stain (note the stain saturated in the white matter). Perls’s stain was used to visualize iron. Primary (V1) and secondary (V2) visual cortices are indicated.

Within the cortex the maps of all quantitative parameters revealed layer-dependent contrasts reflecting the fine details of cortical myeloarchitecture, showing the increase of cortical myelin and cortical iron content from pial surface to white matter boundary and highly myelinated layer IV. Moreover, qMRI maps reflected clear differences in cortical myeloarchitecture between brain areas with different cytoarchitecture as exemplified by the border between primary and secondary visual cortices (**Figure 2, bottom**).

While R2* and MTsat maps demonstrated a homogeneous appearance and a constant gray-white matter contrast across the brains, R1 maps of some specimens exhibited a gradient between the center of the brain and its periphery (**Figure 2, top**). As perfusion fixation is technically not feasible for our implemented tissue sourcing pipeline, this gradient was likely a diffusion fixation artifact.

Thus, MTsat and R2* maps proved as more robust and accurate metrics of whole brain tissue microstructure evaluation across the investigated *post mortem* brains.

### Histological validation of qMRI myelin and iron metrics

In a subset of five animals with a wide range of ages (0.1, 1.75, 6, 12, 16, 30), qMRI metrics were validated by qualitative comparison with histological stains that are regarded as a gold standard for visualizing histological features and validating MRI markers^49^. qMRI metrics of myelin (MTsat) and iron (R2*) obtained in *post mortem* chimpanzee brains demonstrated a high correspondence to the histological stains^49^. Multi-modal histological data, including a Gallyas silver stain for myelin visualization, and a Perls’s stain for iron detection, demonstrated the same features of cortical myeloarchitecture as the qMRI maps (**Figure 2** **bottom, Supplementary** Figure 1). The successful cross-validation is exemplified by the occipital’s lobe stria of Gennari, where the sharp boundary between primary and secondary visual cortex could be identified in all qMRI maps and its location corresponded well to the area boundary determined by histology. Variation in the myelin and iron distribution across cortical layers and cortical regions (occipital, central and frontal) was reflected well by qMRI markers (**Supplementary** Figure 1), demonstrating their suitability to assess cortical architecture in *post mortem* sourced ape brains.

Next, we demonstrated that the area-specific inter-individual variation of qMRI metrics measured in *post mortem* brains of different ages reflects the age-related inter-individual differences in cortical microstructure captured by histology, despite the differences in tissue fixation conditions between brains. We compared qMRI and histological metrics of cortical iron and myelin between the brains of a 2 year old and a 12 year old chimpanzee (Figure 2) in primary and secondary visual cortices.

**Figure 2:**
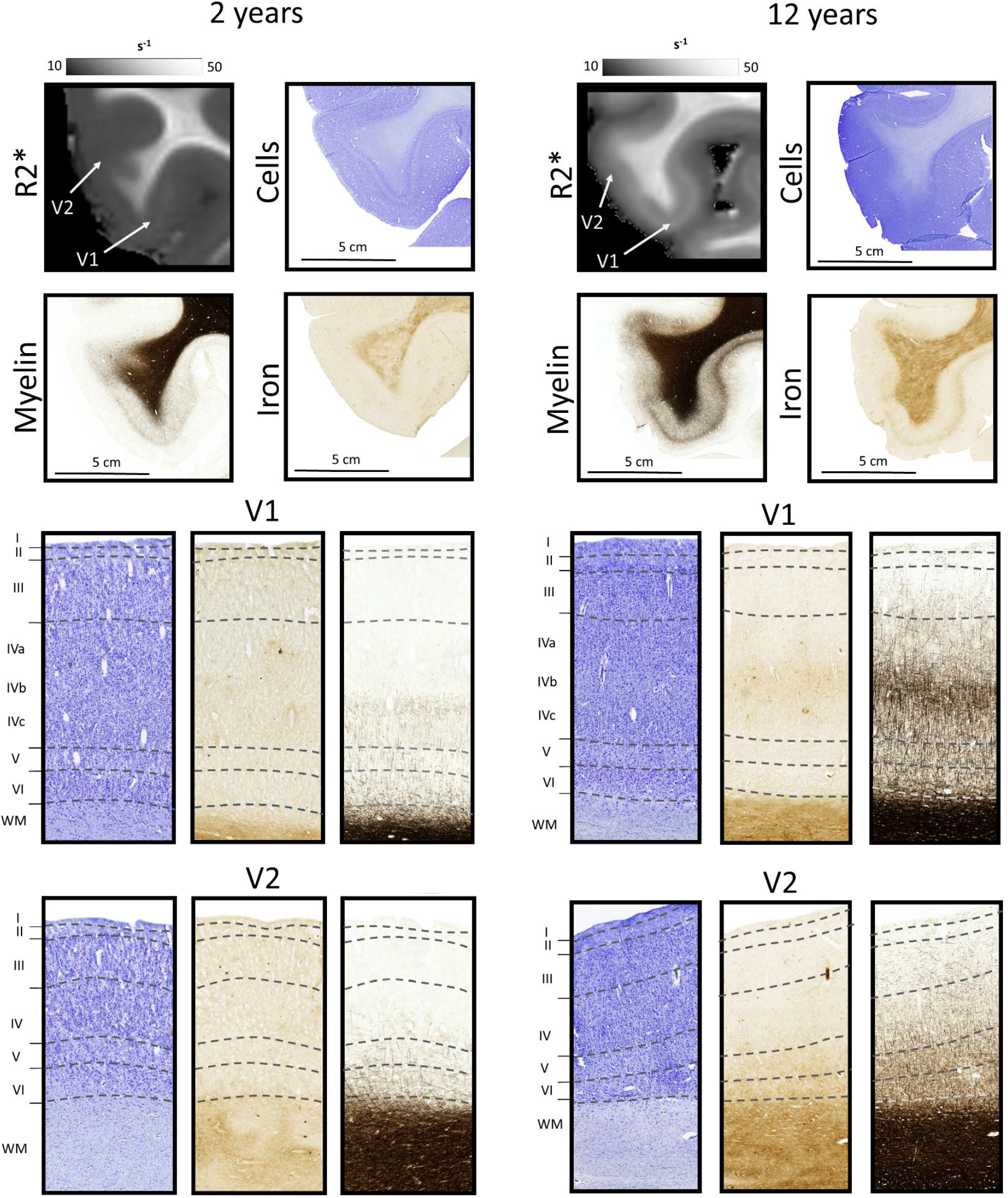
Intracortical laminar contrasts. Histological comparison between the cortex of a wild, 2 year old male chimpanzee’s brain and a sanctuary-housed, 12 year old male chimpanzee’s brain. Cytoarchitecture, iron distribution and myeloarchitecture are shown across cortical depths in sections from primary visual cortex (V1) and secondary visual cortex (V2). Cortical layers are indicated on the Nissl stain. Stronger myelination and intracortical differentiation in both myelin and iron are visible in both V1 and V2 of the 12 year old, compared to the 2 year old. The highly myelinated line of Gennari is observed in V1, particularly for the 12 year old. The primary histological features are well reflected in the R2* maps.

The cortex of the 12 year old was more myelinated, had more iron, and showed more pronounced layer-differentiation in the histology (Figure 2 **and Supplementary** Figure 5). This was reflected by the increased cortical values of R2*, MTsat and by the clearer presentation of the stria of Gennari in the qMRI maps for the older brain. Also the cortical profiles of V1 and V2 in all qMRI maps differed more strongly for the 12 year old’s brain. These additional histological analyses support the use of quantitative MRI for characterizing myelin and iron differences between brain areas, across cortical depth and across age.

### Mapping myelination and iron content across the entire chimpanzee neocortex

Validated ultra-high resolution qMRI was used to map the neocortex of adult chimpanzees. Figure 3 shows the three quantitative MRI parameters sampled at mid-cortical depth together with the maps of cortical thickness for a subgroup composed of all adult chimpanzees (3 females, 6 males, 39.2 *±* 10.5 years old) in the sample. Reconstruction of the pial and white matter surfaces was achieved by careful cortex segmentation. Visual inspection indicates clear differences in cortical myelination and iron content between cortical areas. Primary areas including motor and somatosensory cortices had elevated values, while lower levels of myelination were observed in frontal, parietal and limbic cortices, similar to patterns previously reported on myelin distribution across human brains^33,50,51^.

**Figure 3:**
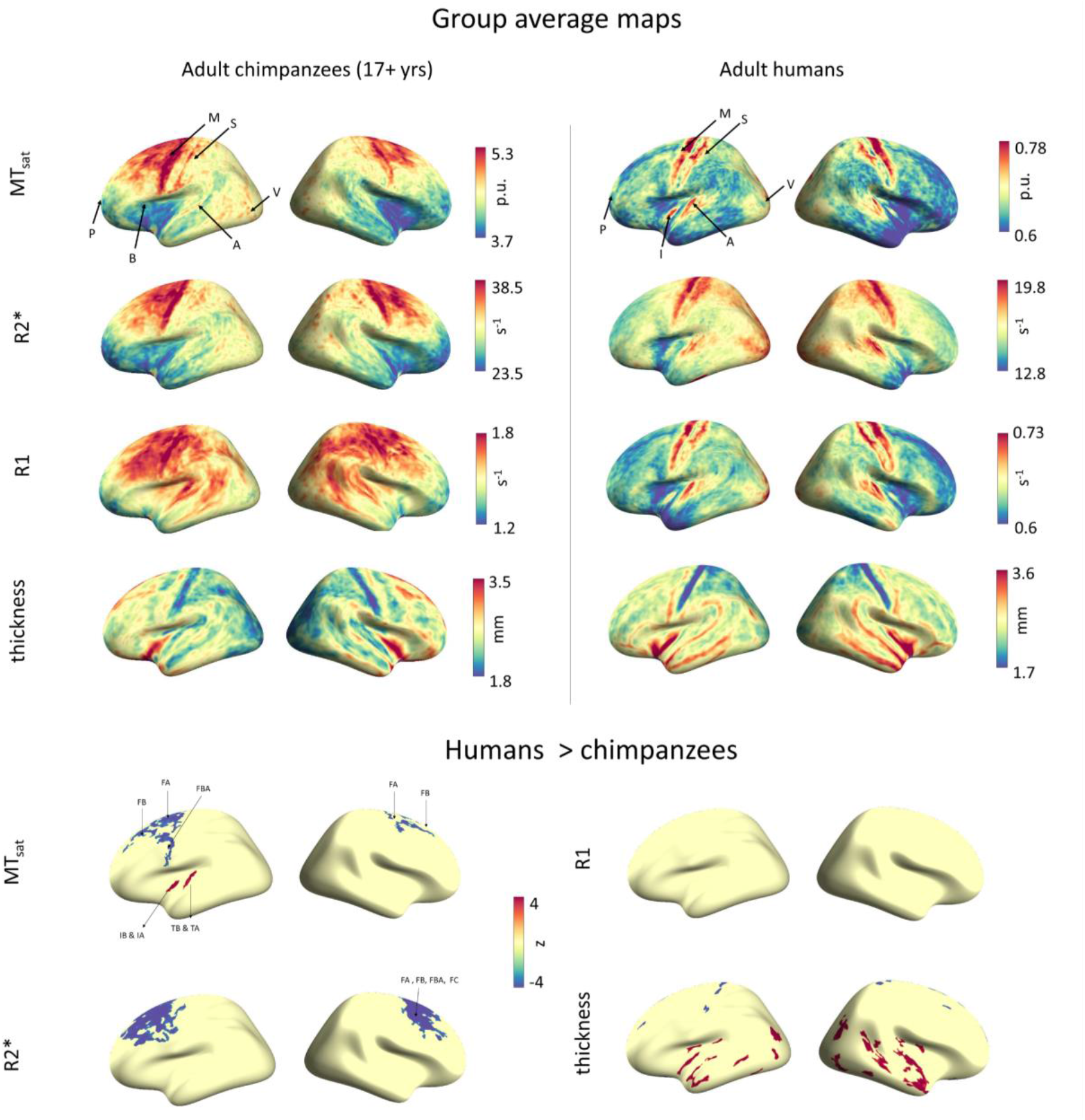
Top: The group-averaged quantitative parameter maps (R2*, MTsat and R1 sampled at mid-cortical depth), and cortical thickness projected onto the inflated surface of the human *fsaverage* template. Data from 9 adult (age > 17 years) chimpanzees and 15 humans contributed to this plot. Display ranges were chosen as 1st and 99th percentile. Noticeable in the chimpanzee brains are the highly myelinated motor cortex (M), somatosensory cortex (S), visual cortex (V), auditory cortex (A) in R1, and the prefrontal cortex (P) and Broca homologue (B) with low myelination. Areas of low cortical thickness include the visual cortices and the postcentral gyrus. In humans, visual cortex (V), auditory cortex (A) and insular cortex (I), areas around the motor cortex (M; including Brodman areas 4 and 6), somatosensory cortex (S; including Brodman areas 1 and 3) stand out as highly myelinated. As a comparison, maps across various cortical depths are shown in **Supplementary** Figure 2. **Bottom:** Statistical comparison between humans and chimpanzees in the cortical distributions of the four parameters. Shown are z-values within statistically significant spatial clusters. A positive z-score (red) indicates higher values in humans than chimpanzees, while a negative z-score (blue) indicates the opposite. Indicated are the BB38 atlas labels FA (primary motor cortex), FB (premotor cortex), FBA (ventral premotor cortex), FC (premotor cortex), IA (anterior insula), IB (posterior insula), TA (superior temporal gyrus), TB (primary auditory cortex). See **Supplementary Table 2** for all significant clusters and their locations.

The visual cortex in the occipital lobe showed slightly elevated values in the MTsat map, but at mid-cortical depths it is not as clearly visible in the R1 map as it is in human *in vivo* data^33,51^. As shown in **Supplementary** Figure 2, the visual cortex stands out more clearly at lower cortical depths.

The maps of cortical thickness demonstrated clear differences between cortical areas (Figure 3). Similarly to humans, the chimpanzee visual cortex is thin and the motor cortex in the precentral gyrus is relatively thick, whereas the adjacent postcentral gyrus is thin.

### Comparison of cortical microstructure between chimpanzees and humans

QMRI metrics in the neocortex were used to compare cortical myelination and iron content between the adult chimpanzee and the adult human brain. QMRI data from 15 healthy human volunteers (age 27.6 *±* 3.8 years; 8 females, 7 males) were analyzed. To statistically compare the myelin and iron maps between species, we computed a vertex-wise two-sample *t*-test. Prior to this, we scaled the chimpanzee maps to match the range of values observed in the humans via a linear model (see **Supplementary** Figure 3, comparable to the data harmonization described in^52^) in order to account for the global differences in the quantitative parameters due to the different scanning conditions (*post mortem* 7T vs *in vivo* 3T). The most striking difference between the two species is the widespread relatively stronger myelination and iron content of the premotor cortex in chimpanzees and the relatively stronger myelination of the auditory cortex in humans, as evident from the statistical comparison of MTsat maps. Additionally, chimpanzees had relatively higher R2* in the motor and premotor cortices. In both humans and chimpanzees, we observed strong spatial agreement between R2* and MTsatt, likely reflecting the colocalization of myelin and iron, due to iron’s role in myelination^29,53^. The only between-species difference in R1 was found in the left medial occipital lobe. See **Supplementary Table 2** for all significant clusters, their size and their location. The between-species differences in the quantitative parameters we found are unlikely to be driven by differences in cortical thickness, which shows a different spatial pattern, with frontal and parietal cortex being thicker in chimpanzees and the temporal cortices being thicker in humans.

### Life span trajectory of the chimpanzee neocortex

The age in our chimpanzee cohort ranged from 3 weeks to 52 years and allowed us to study biological processes through the entire chimpanzee lifespan. The *post mortem* brain specimen studied here also included brains from very young chimpanzees, which are rarely available from captive animals due to their comparatively lower infant mortality.

For each of the brains, we averaged each qMRI parameter across the set of cortical areas, taken from the surface-based cytoarchitectonic Bonin-Bailey (BB38) chimpanzee cortical brain atlas^8^. As cortical segmentation was not possible for the 0.1 year and 1 year old brains due to the minimal gray-white matter contrast, median parameter values across the entire brains were used instead. The age dependence of quantitative parameters was modeled with a simple exponential saturation model. The model was inspired by Hallgren and Sourander’s work on age-related iron accumulation across different brain areas^54^. Our model has three parameters: a constant term describing quantitative MRI parameters at birth; an amplitude, indicating how much the MRI parameter values change from birth to old age; and a rate, indicating how fast the developmental changes are happening. From the rate, a time constant can be derived, indicating how long maturation takes to reach about 60% of the plateau that is reached during adulthood relative to the value at birth.

We performed whole brain analyses. The model parameters (**Supplementary Table 3**) and developmental trajectories (Figure 4) for six particularly relevant regions of interest^26^ are presented in more detail: motor cortex, somatosensory cortex, visual cortex, auditory cortex, frontal pole in the prefrontal cortex, and Broca homologue. For MTsat and R2*, the exponential model explained between 49% and 86% of the variance in the metrics, which was a statistically significant amount in all regions. A previously suggested model describing age-dependence of cortical myelination and iron content by a quadratic polynomial^26,55^ did not explain more variance in the data than our exponential model for any of the regions and all quantitative parameters (**Supplementary Table 3**). The useful property of the exponential saturation model is that it allows one to separately determine the adult level of cortical parameters (i.e. plateau) and the time constant. These two parameters are inherently mixed in the previously reported polynomial models.

**Figure 4:**
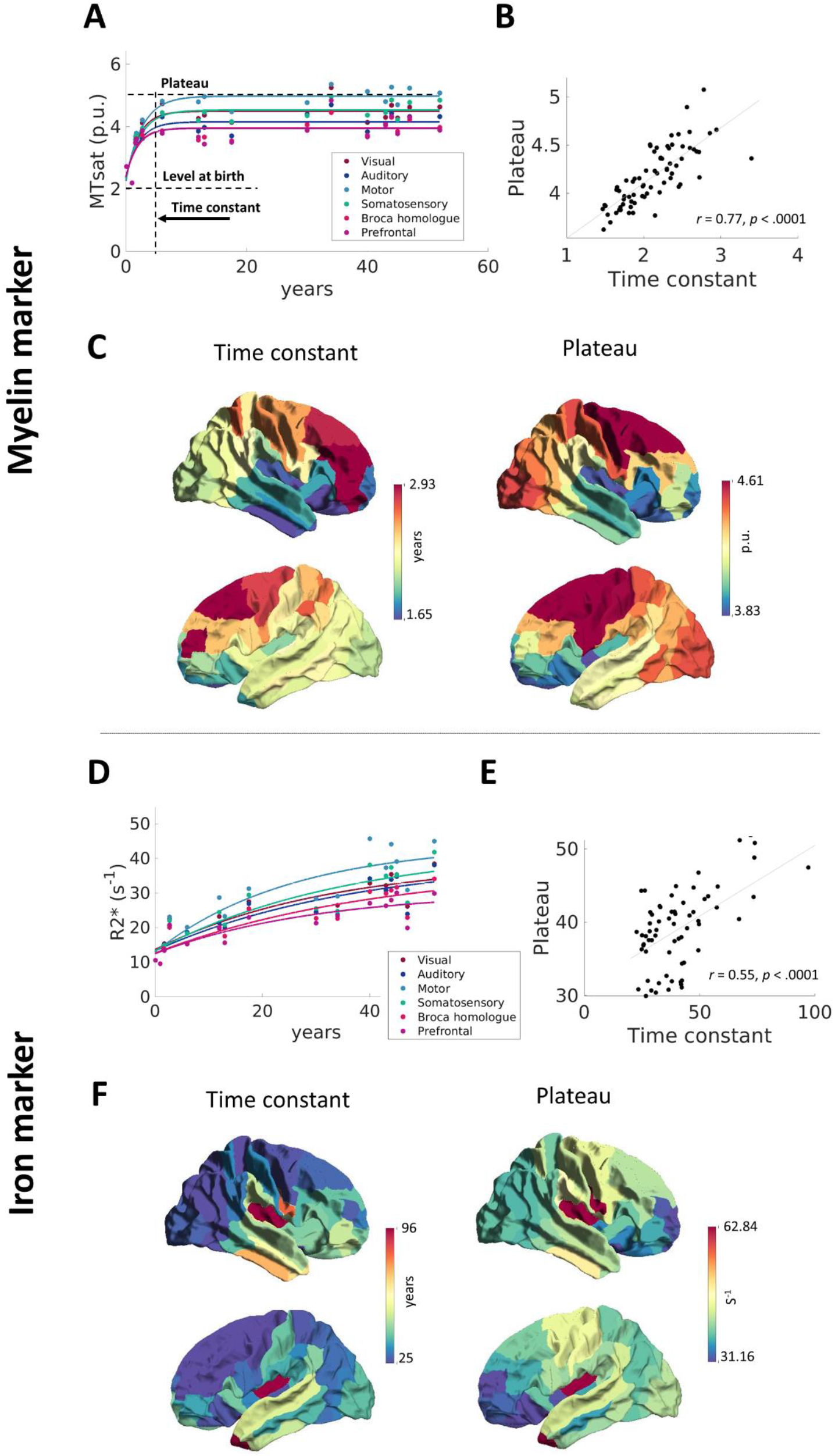
A: For each region of interest, the individual data points and fitted developmental curves for the myelin-sensitive parameter MTsat (left) and the myelin- and iron-sensitive parameter R2* (right) are shown. The model by Hallgren and Sourander^54^ was fit to the data points to illustrate the age trajectories (A and D). Determined are the reached plateau and the time at which around 60% of the plateau has been reached (‘time constant’). The spatial distribution of the parameters for the entire cortex are also shown by assigning each BB38 atlas region a color based on its estimated time constant and plateau, using a colorbar ranging from dark blue (low value) to dark red (high value) (C and F). A higher constant indicates a longer developmental process. The plateau reached during development is also shown. The spatial distributions between time constant and plateau show some correspondence, confirmed by a correlation analysis across all regions between rate and plateau (B and E; Note that regions with no significant model prediction and regions whose value deviated by more than 3 median absolute deviations from the median for either measure were excluded from these calculations). The relationship indicates that more strongly myelinated regions take longer to fully myelinate, and that regions with more iron accumulate over a longer period of time.

For R1, only poor model fits (14-29% of the variance explained) could be achieved in all brain areas, with only the motor cortex showing a significant model performance (**Supplementary Table 3**). The R1 values showed strong inter-individual variability across the adult chimpanzee brains, supporting the observation that R1 is more susceptible to variation in tissue handling and quality, such as fixation^56,57^. Therefore, the following analyses focused on the measures MTsat and R2*.

### Developmental myelination

Age-dependences of cortical myelination quantified by MTsat for the subset of cortical regions are shown in Figure 4A. MTsat^46^ steadily increased and roughly doubled with age in all cortical areas reflecting developmental increase of cortical myelination. The time constant of exponential saturation varied between cortical areas and ranged between 1.5 and 3 years (Figure 4C), with an averaged value of 2.1 years across the entire cortex. Thus, based on our employed model, the myelination of all cortical areas in the chimpanzee brains does plateau and is largely completed by the age of 9 years. Figure 4 also shows the negative correlations between time constant and reached plateau, indicating that areas with higher myelination levels in adulthood take longer until these levels are reached. For example, the highly myelinated motor and premotor cortices had the longest myelination periods. This is in contrast to the often stated assumption that cortical areas high in myelin mature most quickly^19,21,58^.

### Age-related iron accumulation

Age-dependence of iron content in the chimpanzee cortex, quantified by R2*^59^, is shown in Figure 4D. R2* increased from childhood throughout adulthood to old age by around 300%, indicating that iron accumulation continues throughout the lifespan. The time constants for iron accumulation varied between 25 and 40 years and were much higher than that reported for the human brain, where time constants of 20, 16.7 and 14.3 years suggest comparatively fast iron accumulation in motor, occipital and frontal cortex respectively^54^. The regions with higher adult iron levels accumulate less quickly as demonstrated by the negative correlation of plateau and time constant.

### Cortical thinning

Significant decreases in cortical thickness were only found in a subset of cortical regions, including prefrontal, visual, parietal and somatosensory cortices (for all regions see **Supplementary Table 4**), corroborating a previous lower resolution MRI study in captive chimpanzees^10^. We could not find significant thickness changes with our model in the other areas. Thickness changes are comparatively low in comparison to the inter-individual variability in thickness, which makes them difficult to reliably estimate with our sample size.

### Intracortical profiles

Differences in cortical myeloarchitecture are not solely reflected in the myelin concentration sampled at one depth in the cortex but also the pattern of myelination across cortical layers. Early histological work partially captured the chimpanzee myeloarchitecture by studying selected brain sections in individual chimpanzee brains^60–63^. Using our 3D ultra-high resolution data, we have measured average intracortical profiles across the entire brain of a cohort of chimpanzees. To do this, we applied a computational equivolume layering approach (which adjusts surface placement to the local cortical folding^64,65^) to estimate layer positions between the white matter and pial surfaces. We then extracted the quantitative MRI parameters across cortical depths within the individual atlas-defined regions. Figure 5A shows the group-averaged intracortical profiles for the six regions of interest. An increase from pial to white matter surface is clearly visible for all quantitative parameters, reflecting well the expected myeloarchitecture^66^. To emphasize the myeloarchitectonic differentiability of different cortical regions, we regressed out the global profile (average profile across all atlas regions) and show ‘residual profiles’. These show six distinct patterns from pial to white matter surface (Figure 5A), further emphasizing the microstructural differentiation in the chimpanzee brain. This differentiation is also reflected in the spatial maps of a cortical-depth weighted skewness score, which we calculated as described in^67^. The skewness quantifies the extent to which the typical intracortical myelination gradient (more myelin in deep compared to superficial layers) is seen, which is reflected by negative values. Using MTsat as a metric, the most negative average skewness values are found in the visual and motor cortices, and the least negative skewness in frontal cortices (Figure 5B). The spatial distribution is comparable to the one reported for humans in previous work where skewness was quantified without consideration of the cortical depth^24^. Using intracortical profiles based on R2* for the skewness calculation, the least negative values are obtained in visual and motor cortices, and the most negative values in frontal cortices. This indicates that while MTsat and R2* have similar distribution across the cortex, they are sensitive to different aspects of intracortical architecture.

**Figure 5:**
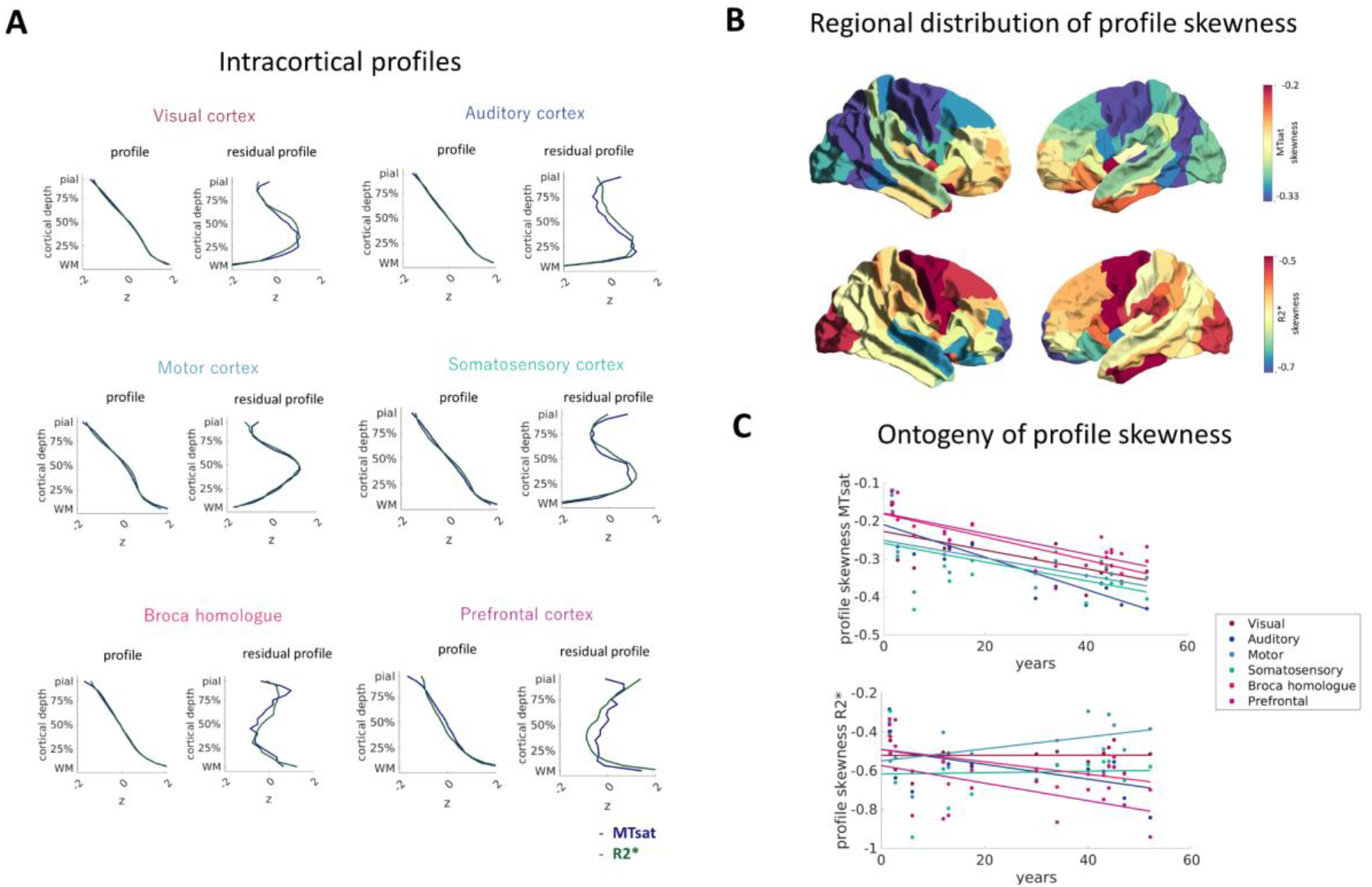
Intracortical profiles. A. Intracortical profiles for six regions of interest are shown as depth- sampled quantitative metrics from pial to white matter surface. The intracortical profile was averaged across hemispheres and chimpanzees after z-standardization (we averaged across both hemispheres, as the profiles were highly reproducible between hemispheres). The whole-brain average profile was regressed out to show region-specific deviations from the whole brain profile, here called ‘residual profile’. Profiles of the parameters MTsat and R2* show similar trends but distinct differences that are reflected in the skewness distributions. B. The profile skewness averaged across chimpanzees and within each atlas region is shown here. Note the particularly negative values in myelin-based (MTsat) skewness for primary motor and primary visual cortices and the least negative values in the prefrontal cortex, comparable to24. Interestingly, the iron-based (R2*) skewness shows a different, inverse, distribution across the cortex. C. For each chimpanzee and region, the skewness of the intracortical profile was quantified. A linear decrease in skewness measures from childhood to adulthood was found for MTsat-based skewness. Supplementary Figure 4 shows average intracortical profiles grouped by the chimpanzee ages for illustration.

### Maturation of intracortical profiles

To study the ontogenetic development of myeloarchitecture and cortical layer-specific maturation, a linear developmental model was applied to the skewness measures. In all regions, the skewness of the MTsat-based profiles was generally negative and decreased with the chimpanzees’ age (Figure 5). This is in line with previous results^24^ in humans. For R2*-based profiles, skewness only significantly decreased in the auditory cortex and Broca homologue (for stats see **Supplementary Table 3**).

## Discussion

This study characterizes the myelin and iron distribution across the entire chimpanzee cortex and across the chimpanzee lifespan using ultra high-resolution ultra-high field quantitative MRI markers. We validated qMRI markers of myelin and iron distributions with gold standard histology on a subset of samples. We used the quantitative character of qMRI markers to compare cortical microstructure between chimpanzees and humans, and to provide developmental trajectories of cortical myelination as well as iron accumulation across the entire chimpanzee lifespan and to determine changes in intracortical profiles across age, reflecting cortical depth-dependent maturation.

To our knowledge, patterns of cortical myelination and iron have not yet been reported for entire chimpanzee brains. Efforts to map the chimpanzee brain have been limited to MRI data obtained from anesthetized captive animals at comparatively low resolution (19 x larger voxel volume), and a single myelin-sensitive contrast, the T1w/T2w ratio^6,41^. T1w/T2w is frequently used as a myelin-sensitive contrast in humans^19^, but the interpretation of this contrast is not straightforward due to technical limitations and focus on a single composite metric^68^. We demonstrated that the inter-regional variation in quantitative MRI goes hand in hand with variation in staining intensities in myelin- and iron-specific histology. Additionally, clear differences in staining intensity and patterns were observed between a 2 year and a 12 year old chimpanzee brain. This histological analysis provides a qualitative addition to the MRI, facilitating the biological interpretation of the quantitative MRI results.

Whole-brain surface projections of the quantitative MRI microstructural biomarkers showed pronounced variation in the quantitative parameters across cortical areas, reflecting the known variability in cortical myelination, including the highly myelinated primary motor, somatosensory, auditory and visual cortices. These cortical myelination patterns complement previous observations in humans and chimpanzees *in vivo*^20,34,41^. While our human cohort was on average slightly younger than the chimpanzee cohort, global age effects are accounted for through our harmonization model. The regional comparison to quantitative maps in humans revealed particularly strong myelination and iron accumulation in motor and premotor cortices in chimpanzees, and stronger myelination of the human auditory cortex.

While the MTsat and R2* maps have similar distribution, probably due to iron’s role in myelination^29^ and the additional sensitivity of R2* to myelination, the difference in specificity of our MRI markers is well demonstrated by the different developmental trajectories between our myelin marker and our iron marker. In all investigated cortical regions, MTsat stayed relatively stable during adulthood, after increasing strongly during youth, likely reflecting developmental myelination^25^. To quantify developmental speed, we applied the exponential saturation model by Hallgren and Sourander^54^ which estimates a rate and corresponding time constant at which a developmental plateau is reached. The time constant is independent of the measurement units and height of the plateau, and therefore provides a basis for comparison within and across species even if not conducted with the exact same methods. The regional variation of the developmental time constant across the cortex suggests a systematic pattern, with slowest myelination of the motor and premotor cortices and fastest myelination in the prefrontal cortex, Broca homologue and auditory cortex. The time constants for myelin lay below 3 years, reflecting the early developmental increase, and a suggested completion of the process by 9 years of age, which is in contrast to the human literature, which suggest that myelination continues until the 30s^14,58^.

We found that cortical areas that have higher levels of myelin in adulthood take longer to complete their myelination. Early histological studies on brain development postulated the opposite relationship. They observed that highly myelinated areas myelinated earlier ^21,69^ and conclude that their myelination is also completed earlier. This fundamental conclusion was recently challenged by both histological studies and particularly by quantitative neuroimaging data in humans, that report a variety of spatio-temporal patterns of cortical and white matter tracts myelination^58,70–72^. This discrepancy may be attributed to the strength of the quantitative MRI, which allows direct quantitative comparison of myelination metrics between brain areas, between individuals and across lifespan. This is prerequisite for precise modeling of developmental brain trajectories, which allows to disentangle the speed of myelination and the time of its completion.

One previous study assessed myelin development in a group of 20 chimpanzees in comparison to 33 humans through histological quantification of cortical myelination using histology-based stereology^26^. Age-related increases of myelin were reported for visual cortex, motor cortex, somatosensory cortex and frontopolar cortex in both species. However, fitting of different polynomial developmental trajectories indicated a protracted myelination in humans compared to chimpanzees. Since the study used different histological methods for the human and chimpanzee tissue, pooled data across the adult chimpanzee, and the model confounded the speed of myelination with its strength, it complicated the interpretation of the results. Additionally, the quadratic model that was fit to the data indicates myelin decline pre-adulthood, which would be difficult to explain neurobiologically^14^. Comparing the model fits in our data, their quadratic model for development until the age of seventeen^26^ explained less variance than our exponential model, indicating that the trajectory of developmental myelination is better described by our approach.

In contrast to MTsat, R2* steadily increased from birth and continued to increase into adulthood. This effect has also been shown in humans *in vivo*^35^ and is most likely due to its sensitivity to iron, which accumulates with age. Iron accumulation is also slowest in the motor and premotor cortices and faster in prefrontal and temporal cortices. The time constants suggest a slower iron accumulation in chimpanzees than what was reported by Hallgren and Sourander^54^ in human cortices. Our results indicate that cortical iron accumulation, a process related to neurodegeneration in human aging^30^, is also present in chimpanzees. One reason why humans are more susceptible to neurodegenerative disease may be this faster iron accumulation and the generally longer lifespan that also leads to the inverted U-shape trajectory of myelination found in imaging studies. This shape is driven by an increase of myelin into early adulthood and a decline from age 60^73^, both in white matter^35,39^ as well as in cortex^58,74,75^. We did not observe this U-shaped trajectory in our chimpanzee data, and one reason may be that the oldest chimpanzee in our sample was 52 years old, reflecting the generally longer lifespan in humans and the challenges in comparing aging processes between the species^9,76,77^.

The ultra-high-resolution quantitative MRI revealed strong intra-cortical variation in myelination. In particular, area-specific myeloarchitectonic features like the highly myelinated stria of Gennari in the primary visual cortex, and borders between areas with known cyto- and myeloarchitectonic differences (such as primary and secondary visual cortex) were clearly visible. Intracortical profiles obtained from various regions defined by the histology-based cortical Bonin-Bailey chimpanzee atlas clearly differed from each other, which was also reflected in the derived skewness measures. Negative skewness indicates preferentially stronger myelination in middle to deep layers ^24,67^. In line with this, we found the strongest negative skewness in the motor cortex, followed by visual and auditory cortex, with the weakest negative skewness in the Broca homologue and frontal pole. The distribution of skewness across the cortex corresponds well with a previous study in humans^24^. The R2*-based skewness distribution showed the opposite trend, potentially providing a complementary histological measure.

We also explored intracortical ontogenetic development, which has been studied in humans in previous histological work^78^, and recently *in vivo* with MRI^24^ which showed that the cortical development is not only reflected in an increase in myelin-sensitive parameters, but also in a decrease in the skewness of the intracortical profiles, which is likely a result of preferential developmental increases in myelin in mid- to deep cortical layers. In our data, skewness in the MTsat-based profiles also decreased with the chimpanzee development from childhood to adulthood.

The MRI findings suggest that our whole brain ultra-high resolution approach provides a feasible addition to histological studies on cortical microstructure^60–63^. In contrast to previous studies in chimpanzees, our imaging approach has the following advantages: its multi-modality provides three different quantitative parameters that differ in their sensitivity to various tissue characteristics^36,37^, which allowed us to study different biological aspects of cortical maturation. It captures whole chimpanzee brains, whilst histological studies rely on few specimens and on small regions or are limited to specific cutting planes and are not quantitative in nature. The creation of population-based cortical parcellations for chimpanzees could therefore in the future be achieved with our MRI approach that covers different microstructural aspects. QMRI is also the method of choice used in human developmental studies^24,25^, facilitating between-species comparison through literature comparison or the combination and harmonization of datasets.

Several caveats are worth highlighting. While our sample size is comparable to previous studies^26^, it was not sufficiently large to be able to test and select from a variety of possible developmental models, or to statistically compare model parameters across cortical regions. In some atlas regions, the model did not perform well (see **Supplementary Table 4** for modeling parameters for all BB38 atlas regions).

The employed qMRI parameters are indirect markers of myelination and iron that may be influenced by different factors, especially in case of pathologies^36^. However, the careful histological validation within the specific setting of this lifespan study indicates that the qMRI parameters can be regarded as reliable microstructural markers.

The comparison of chimpanzee and human lifespan changes required the harmonization of the *post mortem* chimpanzee and *in vivo* human data, in order to compensate for differences due to the different data acquisition conditions.

Our tissue collection strategy implies some uncontrollable variation in *post mortem* intervals and fixation conditions between sourced brains, which may influence the observed inter-individual differences in the quantitative parameters. MRI data suggested high tissue quality in most brains despite sometimes challenging tissue collection conditions in African field sites. Data from three individuals had to be excluded to severe MRI-visible cortical pathologies, induced by systemic bacterial infections causative for the animals’ death. Additionally, in three brains, fixation-related artifacts were observed in the R1 maps only, but not in the other parameters. R1 seemed to be the most prone to artifacts that are probably related to fixation. This may have led to pronounced inter-individual variability which obscured the developmental trajectory. Previous work has confirmed a link between fixation protocol and quantitative MRI parameters, in particular for R1^57^. While we kept the fixation protocol as consistent as possible, some factors still varied between the brains, such as the interval between time of death until fixation, constant fixation temperatures and non-standardized formaldehyde solutions purchased in the country of brain sourcing. This highlights the advantage of multi-parametric mapping, where several metrics are obtained that capture different physical properties of the tissue.

## Conclusion

We showed qMRI-based myelin and iron distribution across the entire cortex in chimpanzee brains and differences to humans in motor and premotor cortices. We demonstrated steep changes in myelination from birth to early adulthood and continuous iron accumulation from birth to old age in chimpanzees. Making use of our ultra-high resolution qMRI data, we also showed that intracortical profiles differed across cortical areas and matured across the chimpanzee life-span.

## Online Methods

### Brains

The brains used in this study were obtained through the Evolution of Brain Connectivity Project following its associated standard procedures^42^. Brains from wild and captive chimpanzees (Pan troglodytes) were sourced from African field sites, zoos and sanctuaries. The brains were extracted within 4-24 hours after death, and fixed for 8-12 weeks in 10 % phosphate-buffered saline (PBS) buffered formalin. The samples were then shipped to Leipzig where superficial blood vessels were removed and the formalin was washed out in PBS with 0.1 % sodium azide at pH 7.4. The procedures were in line with the ethical guidelines of primatological research at the Max Planck Institute for Evolutionary Anthropology, Leipzig, which were approved by the ethics committee of the Max Planck Society.

20 chimpanzee brains were included in this study, 5 of them for histology. An overview of the brains is provided in Supplementary Table 1. Three brains were excluded from all analyses due to low tissue quality, yielding usable data from 17 brains.

The causes of death for chimpanzees from field sites included bacterial infections (N = 4), conspecific aggression (N = 3), human-animal conflict (N = 2), leopard attack (N = 2), emaciation due to chronic renal disease (N = 1) and emaciation due to maternal loss (N = 1). The cause of death for chimpanzees that died at sanctuaries and zoos were cardio-vascular diseases (N = 1), tumors (N = 1), conspecific aggression (N = 2), renal disease (N = 1), epileptic episode (N = 1), and unknown causes (1).

For 9 chimpanzees who had truncated life history data (immigrant females, individuals that were present at the start of the observation, unhabituated individuals) we estimated the age according to behaviour, size, sexual and dominance activity, face and body appearance in relation to known sex-age-class relationship, with an error margin related to how exact this estimate is (0-10 years: ± 1 year, 10-15 years: ± 3 years, >15 years: ± 5 years; see^44,45^).

### MRI acquisition

For scanning, the whole brains were immersed in the proton-free solution Fomblin (Solvay) in a tightly sealed plastic container. Data were acquired on a human whole-body 7T Terra MRI scanner (Siemens Healthineers, Erlangen, Germany), using a 32 channel receive, 1 channel transmit radio- frequency (RF) human head coil (Nova Medical, Wilmington, MA). The brains were kept at room temperature for at least 2 hours before the start of the scanning. To control for the temperature dependence of quantitative MRI parameters and minimize heating effects during imaging, the sample temperature was kept at 30°C with a precision of ±2° using warm air flow. The temperature was monitored throughout the scanning session using a fiber optic temperature sensor (LumaSense) attached to the outside of the scanning container, using the TrueTemp software (Luxtron).

Multi-parametric mapping was implemented using a multi-echo 3D FLASH sequence^79,80^ at 300 μm isotropic resolution (matrix: 432 x 378 x 288; readout bandwidth of 331 Hz/pixel; repetition time (TR) = 70 ms; 12 equidistant echoes with echo times (TE) between 3.63 ms and 41.7 ms with a bipolar readout, △TE = 3.56 ms; excitation flip angles: 18° (PD-weighted and MT-weighted), 84° (T1-weighted); MT pulse characteristics: Gaussian at 3 kHz offset, flip angle: 700°). The order of scan sequences was kept consistent across scanning sessions. To maximize image quality, no k-space undersampling or other acceleration methods (such as partial Fourier or parallel imaging) were used. The acquisition time for each of the weighted images was just over 2 hours.

The RF receive bias (B1^-^) field was partly corrected using the ratio of a low resolution (2.1 mm) T1- weighted image acquired with the 32-channel receive coil vs an image acquired with the transmit coil in receive mode.

A spin and stimulated echo measurement sequence was used to obtain maps of the RF transmit field B1^+^ and a dual gradient echo sequence was used to map the static magnetic field B0^81,82^.B1^+^ mapping was done at an isotropic resolution of 4 mm, with a matrix size of 48 x 64 x 48 and the following acquisition parameters: TR = 500 ms; TE = 40.54 ms; mixing time TM = 34.91 ms; flip angles 330° to 120° in steps of 15°; RF duration 24 μs per °; GRAPPA acceleration factor = 2 x 2. B0 mapping was done at an isotropic resolution of 2 mm and a matrix size of 96 x 96 x 64, with the following acquisition parameters: TR = 1020 ms, TE = 10 and 11.02 ms, flip angle = 20°. The B0 map was used to correct the EPI distortions in the B1^+^ map.

### MRI acquisition - human data

We used multi-parametric mapping data from 15 healthy volunteers (mean age 27.6 ± 3.8 years (range: 21-30 years), 7 males, 8 females) acquired on a 3T Connectom scanner equipped with a 32- channel receive radiofrequency (RF) head coil (Siemens Healthineers, Erlangen, Germany) and a body transmit coil. Multi-parametric mapping was implemented using a multi-echo 3D FLASH sequence^79,80,83^ at 800 μm isotropic resolution (matrix: 320 x 280 x 224; readout bandwidth of 488 Hz/pixel; TR = 25 ms; 8 equidistant echoes for PD-weighted and T1-weighted images and 6 equidistant echoes for MT-weighted images, with TEs between 2.4ms and 18.64 ms (14 ms for MT-weighted), △TE = 2.32 ms; excitation flip angles: 6° (PD-weighted and MT-weighted), 21° (T1- weighted); MT pulse characteristics: Gaussian at 2 kHz offset, flip angle: 180°, 4 ms). Additional parameters: GRAPPA acceleration factor = 2 × 2. The acquisition time for each of the weighted images was about 7 min.

B1^+^ mapping was done at an isotropic resolution of 4 mm, with a matrix size of 64 x 64 x 48 and the following acquisition parameters: TR = 500 ms; TE = 43.5 ms; mixing time TM = 38.2 ms; flip angles 230° to 130° in steps of 10°; RF duration 28 μs per °; GRAPPA acceleration factor = 2 x 2.

B0 mapping was done at a resolution of 3 x 3 x 2 mm and a matrix size of 64 x 64 x 64, with the following acquisition parameters: TR = 1020 ms, TE = 10 and 12.46 ms, flip angle = 90°. The B0 map was used to correct the EPI distortions in the B1^+^ map.

The RF receive bias (B1^-^) field was partly corrected using the ratio of a low resolution (4 mm) PD-weighted image (excitation flip angle 6°, TR 6 ms, TE 2.4 ms) acquired with the 32-channel RF receive coil vs an the image acquired with the body RF coil in receive mode. Separate receive bias field maps were acquired for the PD-, T1- and MT-weighted acquisitions to account for the inter-scan motion effects on the bias field^84^.

The study was presented to and approved by the ethics committee of the medical faculty of the University of Leipzig (ID: 293/18-ek). All subjects gave written informed consent before being scanned.

### MRI analysis

#### Calculation of the quantitative MRI maps – post mortem data

The B1^+^ maps were computed with the open source hMRI toolbox^85^ (https://hmri.info), using an approximate global reference T1 of 500 ms (which accounts for T1 recovery during the mixing time; this is shorter than the default in the toolbox due to the shorter T1 of fixed tissue compared to *in vivo* T1). The spatial smoothing of the maps was adjusted for our data, which are characterized by sharp edges between the brain and the no-signal background. We implemented a boundary-preserving smoothing procedure: a median filter of an 8 mm Gaussian smoothing kernel was applied to the B1^+^ map and then divided by a brain mask (which we obtained through an intensity thresholding of the first echo of the T1w images) that had been smoothed in the same way. The resulting B1^+^ maps were then upsampled to the high resolution images. A similar processing strategy was used for the B1^-^ images, which were calculated by dividing the image intensity obtained with the 32-channel receive coil by the acquisition with the transmit coil in receive mode (done for the first five echoes and then averaged).

All weighted FLASH images were corrected for off-resonance-related distortions in the readout direction, which alternated between odd and even echoes due to the bipolar readout scheme. First the geometric mean of the first and third echo was calculated, which is an estimate of a virtual second echo image acquired with the opposite readout polarity^86^. Using the second acquired echo and the virtual echo as input, the HySCO algorithm of the ACID toolbox (http://diffusiontools.com/) was used to estimate the distortions and correct all acquired echoes^87,88^.

For the analysis of one of the 17 brains we had to correct for movement of the brain in the container over the duration of the experiment. To do this, we added an additional first step in the pipeline, where the three weighted images were first registered, by finding an affine mapping of the first echoes to the first echo of the T1w image (using FSL flirt with 12 degrees of freedom). No registration between contrasts was done for the other brains.

The effective transverse relaxation rate (R2*) was then fit using weighted least squares, as implemented in the hMRI toolbox^85,89^. From this fit, the weighted images were extrapolated to TE =These images were then used to calculate R1, PD and MTsat as described in^43^. The MTsat images were additionally corrected for residual B1^+^ bias using the global parameter from^43^.

#### Calculation of the quantitative MRI maps – human data

B1^+^ maps were computed using a global reference T1 of 1192 ms, the default 3T *in vivo* value specified in the hMRI toolbox. As for the *post mortem* chimpanzee MRI data, the effective transverse relaxation rate (R2*) was then fit using weighted least squares and the weighted images were extrapolated to TE = 0. These images were then used to calculate MTsat, R1 and PD maps, as implemented in the hMRI toolbox. The map creation also included a per-contrast RF sensitivity bias field correction as well as a correction for imperfect spoiling.

#### Cortex segmentation

As the scanning was done in an “MR-invisible” proton-free solution, we obtained brain masks by applying an arbitrary intensity threshold to the T1-weighted images obtained at the shortest echo time. For tissue segmentation, we downsampled the MTsat images, which provided the best contrast between gray and white matter, to a resolution of 0.7 mm isotropic. We then used a pipeline of functions from the software package Freesurfer (https://surfer.nmr.mgh.harvard.edu^90^) with the following steps. steps of the *recon-all* pipeline for segmentation of the cortex and surface reconstruction: *autorecon1* was applied (flags: -noskullstrip -notal-check -hires), the brain mask (calculated as described above) was converted to Freesurfer format with *mri_convert* (flag: -- conform_min), and autorecon2 was run (flags: -notal-check -hires). We then manually edited the white matter masks and/or the contrast in the *brain.mgz* image as necessary in *freeview*, and reran the Freesurfer segmentation with *autorecon2-wm* (flags: -notal-check -hires). Pial surfaces were refined with *autorecon-pial*, and surface-based registration (to the human Freesurfer template) was achieved with *autorecon3* (flags: -notal-check -hires).

Cortical surfaces for the *in vivo* human data were again reconstructed using the *recon-all* pipeline from Freesurfer. As the contrast in qMRI maps deviates significantly from the T1w MPRAGE image contrast expected by the *recon-all* pipeline^83^, the following steps were taken to extract an image with MPRAGE-like contrast from the 3T qMRI parameter maps^91^. First, a small number of negative and very high values produced by estimation errors were removed from the R1 and PD maps. Then, these two maps (PD and T1 (= 1/R1)) were used as input to the FreeSurfer *mri_synthesize* routine to create a synthetic FLASH volume with optimal white matter (WM)/gray matter (GM) contrast (TR 20 ms, FA 30°, TE 2.5 ms).

This synthetic image was then used as input to *autorecon1* (flags: -noskullstrip). Skull-stripping was performed by applying a combined GM/WM/cerebrospinal fluid (CSF) brain mask (threshold: tissue probability > 0) with tissue probabilities for the *autorecon1* output image calculated by SPM_segment (https://www.fil.ion.ucl.ac.uk/spm). The skull-stripped synthetic image was then used as input for the remaining steps of the recon-all pipeline.

#### Parcellation

Freesurfer’s *mris_ca_label* was then used to perform surface based registration of the Bonin-Bailey (BB38) cortical brain atlas to the individual chimpanzee’s surfaces^8^. The following regions of interest were defined and parcellated using the BB38 atlas: Visual cortex (BB38 label: OC, corresponds to Brodmann Area 17), Auditory cortex (BB38 label: TB, corresponds to Brodmann Area 41), Motor cortex (BB38 label: FA), Somatosensory cortex (BB38 label: PC, corresponding to Brodmann Area 2), Broca homologue (BB38 label: FCBm, corresponding to the posterior inferior frontal gyrus), Frontal cortex (BB38 label: FDm, corresponding to Brodmann area 9).

For two of the brains (0.1 years and 1 year old), the low contrast between gray and white matter, which is driven by the low myelination at this developmental stage, did not allow for a segmentation with Freesurfer. For these two brains, median parameter values across the entire brain were used for developmental analysis.

#### Intracortical profiles

We applied an equivolume layering approach which adjusts surface placement to the local cortical folding^64,65^ (https://github.com/kwagstyl/surface_tools) to our white matter and pial surfaces, across 5-95% of the cortical depth in 5% steps. The multiparametric maps were then projected onto these surfaces. For each ROI, the median value (R2*, R1, MTsat) across all vertices of each ROI’s layer was calculated. From these values, intracortical profiles were constructed. To obtain an overall measure of regional maturation, the median value (R2*, R1, MTsat) across all cortical depths was calculated.

For averaging cortical profiles across all different individuals, a z-standardization for each profile was performed first and then the median and interquartile range were calculated. Additionally, to illustrate developmental effects, median (not standardized) profiles were calculated for young (<6 years, preadult < 17 years and adult chimpanzees). Robust metrics were chosen to be robust to outliers (profiles of the 1.6 and 1.75 year old chimpanzees often stood out).

To emphasize the subtle myeloarchitectonic differences, we also calculated ‘residual profiles’: here, the average profile across all regions from the Bonin-Bailey (BB38) atlas was regressed out of the individual profiles. Residual profiles emphasize the deviation from the globally observed trend of increasing myelination and iron content going from the pial to the white matter surface.

For each individual’s profile in each region of interest, a cortical-depth weighted skewness score was calculated as described in^67^ (also see^24^). The skewness quantifies the extent to which the typical intracortical myelination gradient (more myelin in deep compared to superficial layers) is seen.

#### Surface projections

To project the high-resolution quantitative parameters to the individual surfaces, we first converted them to Freesurfer space (registration of the high resolution quantitative maps into Freesurfer space with *tkregister2* (flags --regheader --noedit) to obtain the registration matrix; followed by *mri_vol2vol* (flag --no-resample) to apply the registration matrix). The metrics sampled at mid-cortical depth were then projected to the mid-cortical surface from the equivolume layering, using *mri_vol2surf* .

#### Surface-based group statistics

The surface projections were corrected for vertex-wise outliers that could affect group statistics in individual locations. To do this, outliers in each surface projection file were identified as data points deviating more than 3 scaled mean absolute deviations away from the median, using Matlab’s *isoutlier* command, and consequently replaced by the identified lower and upper thresholds of the remaining distribution.

The surface projections of the quantitative parameters were concatenated across all individuals, using *mris_preproc*, followed by a smoothing step (3mm full-width - half-max kernel). Average maps were computed using *glm_fit*.

For the comparison between chimpanzees and humans, we first accounted for global differences in the parameters due to the different scanning conditions (*in vivo* and *post mortem*). To do this, we calculated group means and standard deviations within all BB38 atlas regions for both humans and chimpanzees and then fitted a weighted total least-squares linear model^92^ across the 76 regions to obtain a global model for translating the *post mortem* chimpanzee data to the range of values observed in the human *in vivo* data. The model that was obtained for each of the three quantitative MRI metrics based on the group data was then applied to each individual chimpanzee quantitative map. A vertex-wise statistical comparison of the chimpanzee and human maps was obtained through a two-sample t-test using Freesurfer’s *mri_glmfit* and applying a cluster-based thresholding using *mri_glmfit-sim*, with a cluster forming threshold of 3 (p < .001) and a cluster threshold of p < .05 across both hemispheres^93^.

#### ROI-based group statistics

The model by Hallgren^54^ was fit to the data points as MRI-parameter = *a* [1 − exp(−*b* × age)] + *c*, with three parameters to be modeled: a constant term *c* describing quantitative MRI parameters at birth, a term *a* indicating the amplitude of developmental changes, and a rate *b*, indicating how quickly the plateau (sum of constant and amplitude) is reached (the time constant 1/*b* reflects the age at which approximately 63.21% (1/e) of the plateau is reached).

The fitting was done using nonlinear least-squares regression via Matlab’s (R2021a) *lsqcurvefit* algorithm with parameter initializations of 5 (*a*), 0.1 (*b*) and 0 (*c*), a non-negative constraint for all parameters and the default parameters otherwise. The initialization is required for the algorithm to run, but the exact parameters did not affect the outcome. Confidence intervals were calculated through the Jacobian and residuals, using the function *nlparci*. To assess whether the model significantly predicted the data, the predicted MRI parameters (obtained from inserting the respective ages in the estimated model) were correlated with the actual MRI parameters. Only in cases where *p* < 0.05 age effects were deemed relevant.

As thickness is expected to decrease with age, an exponential decrease was fit instead: MRI- parameter = *a* exp(−*b* × age) + *c*, also using a non-negative constraint for *c* and *b*, and a maximum allowed value of 5mm for *c* and *a*.

Age effects of skewness measures were fit with a linear model with no constraints.

### Histology

Brains were cut in coronal slabs and placed in 30% sucrose in phosphate buffered saline pH 7.4 (PBS) with 0.1% sodium azide until cryoprotection was achieved. Slabs of three regions (frontal, central and occipital) from both hemispheres were cryosectioned at a thickness of 30 µm on a cryomicrotome (Thermo Scientific, Microm HM430, freezing unit Thermo Scientific, Microm KS34). The sections were collected in PBS with 0.1% sodium azide and stored at 4°C until further use. Block Face Images were acquired during sectioning. Consecutive sections were stained with histological stains.

*Nissl staining.* To investigate cortical cytoarchitecture, Nissl staining was applied. Sections were mounted on glass slides coated with Poly-L-Lysin, air dried, rehydrated and fixed in 96% ethanol for 30 min, stained with 0.1% acetate buffered cresyl violet, washed in distilled water and differentiated in graded ethanol steps.

*Iron staining.* For visualization of iron distribution, Perls’s stain with diaminobenzidine enhancement was performed^94,95^. Sections were mounted on glass slides, air dried, rehydrated, incubated in the working solution at 37°C and developing reactions carried out under visual control.

*Myelin stainings.* For the silver staining, sections were mounted on glass slides, air dried, rehydrated and processed following the classical Gallyas protocol^96^ with modifications^97^. Finally, sections were dehydrated over graded ethanol steps and embedded with Entellan (Merck) in toluene.

Microscopic slides were imaged with an Axioscan Z1 slide scanner.

## Data and code availability

Data used in this manuscript will be shared upon request. As some primate sites restrict data access for institutions involved in invasive primate research, a data sharing committee will coordinate any data access and collaboration requests and ensure that ethical and permit requirements of sourcing organizations and countries are adhered to.

The pipeline for quantitative MRI data processing and calculation can be found here: https://github.com/IlonaLipp/postmortembrain-mpm

Further analysis scripts for this paper are found in https://github.com/IlonaLipp/LifespanTrajectoryChimpanzees

## Supporting information

All Supplementary Material

## Acknowledgements

This project has been funded by the Max Planck Society under the inter-institutional funds of the president for the Evolution of Brain Connectivity Project and by the European Research Council under the European Union’s Seventh Framework Programme (FP7/2007-2013) / ERC grant agreement number 616905. Nikolaus Weiskopf received funding from the European Union’s Horizon 2020 research and innovation programme under the grant agreement number 681094. This project received funding from the Deutsche Forschungsgemeinschaft (DFG, German Research Foundation)—project no. 347592254 (WE 5046/4-2 and KI 1337/2-2). Ilona Lipp has been funded by the Deutsche Forschungsgemeinschaft (DFG, German Research Foundation)—project number 446291874. Gunther Helms was funded in part by the Swedish Research Council, grant NT 2014-6193. We are grateful to Caroline Jantzen, Felix Büttner, Niklas Alsleben and Franziska Zahn for their help with sample preparation and scanning and segmentation, and to Lenka Vaculčiaková for help with setting up the scan protocol.

We thank the Ministère de l’Enseignement Supérieur et de la Recherche Scientifique and the Office Ivoirien des Parcs et Réserves for permitting the study in Cote d’Ivoire, Uganda Wildlife Authority and Ugandan National Council for Science and Technology for permitting the study in Uganda. Thanks to the staff of the Tai Chimpanzee Project and Budongo Conservation Field Station, Kolmarden Zoo, Chester Zoo and Twycross Zoo for their commitment.

Special thanks to Christina Kompo for her administrative and logistical support. We thank Casey Paquola for her input on the skewness calculation.

