## Supplementary material for "Lifespan trajectory of chimpanzee brains characterized by magnetic resonance imaging histology": All Supplementary Material

**Supplementary Table 1**: Chimpanzee characteristics. Name, age (*Exact date of birth is known; for other individuals it has been determined with ±1 year for individuals under 10 years old, ±3 years for individuals between 10 and 15 years of age, and ±5 years for individuals > 20 years old[44,45]), sex (f = female, m = male), chimpanzee subspecies and acquisition site, are provided. The suspected cause of death and postmortem interval (PMI) are also reported. For each quantitative MRI contrast, it is indicated whether data passed quality check (+) or not (-).

| # | Age (years) | Sex | Subspecies | Site | PMI (hrs) | R1 | MTsat | R2* |
|---|---|---|---|---|---|---|---|---|
| 01 | 6* | f | verus | Field site | 4 | + | + | + |
| 02 | 0.1* | m | schweinfurthii | Field site | 4 | + | + | + |
| 04 | 34* | m | verus | Zoo | 4-6 | - | + | + |
| 05 | 1.75* | m | verus | Field site | 24 | + | + | + |
| 06 | 14 | f | verus | Field site | 24 | - | - | - |
| 07 | 47* | f | verus | Zoo | 3.5-4 | + | + | + |
| 08 | 30 | m | schweinfurthii | Field site | 12 | + | + | + |
| 09 | 13 | f | schweinfurthii | Field site | 15 | + | + | + |
| 11 | 1.6* | m | verus | Sanctuary | <16 | + | + | + |
| 12 | 1 | m | troglodytes | Field site | 8 | + | + | + |
| 14 | 40 | f | verus | Field site | <24 | + | + | + |
| 15 | 2.75* | m | verus | Field site | 3.5 | + | + | + |
| 16 | 44* | f | hybrid | Zoo | 1 | + | + | + |
| 18 | 43 | m | schweinfurthii | Zoo | 1 | + | + | + |
| 25 | 16* | m | verus | Field site | <16 | - | - | - |
| 26 | 15* | m | verus | Field site | <12 | - | - | - |
| 27 | 45 | m | verus | Field site | <18 | + | + | + |
| 29 | 52* | m | verus | Zoo | <24 | + | + | + |
| 32 | 12 | m | verus | Sanctuary | 6 | - | + | + |
| 33 | 17.5 | m | troglodytes | Field site | 11 | - | + | + |

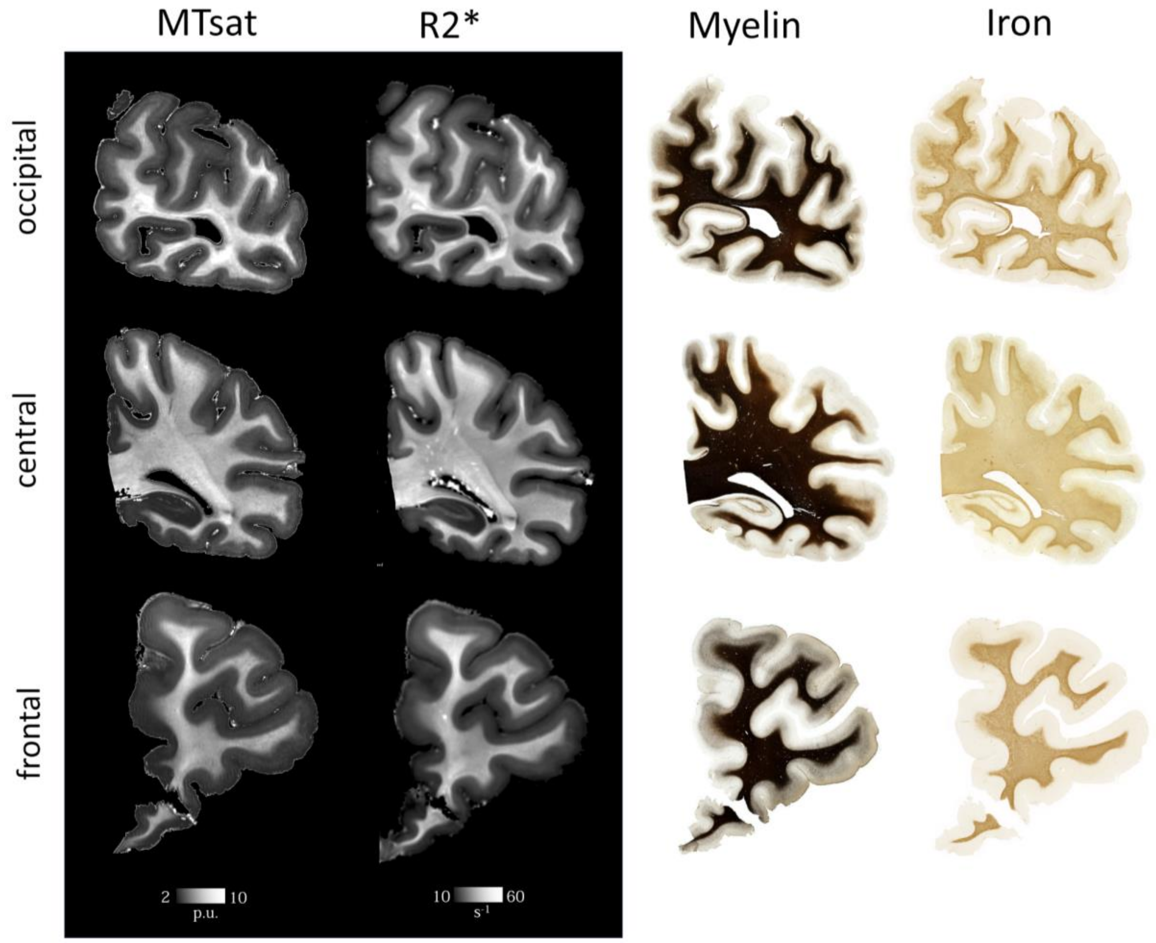

**Supplementary Figure 1**: Quantitative MRI parameters MTsat (myelin-sensitive) and R2* (myelin- and iron-sensitive) and histological stainings of coronal sections of the occipital (top), central (middle) and prefrontal (bottom) lobes in a 12 year old male chimpanzee who died in a sanctuary. Myelin was visualized with a Gallyas silver stain (note the stain saturated in the white matter). A Perls's stain was used to visualize iron.

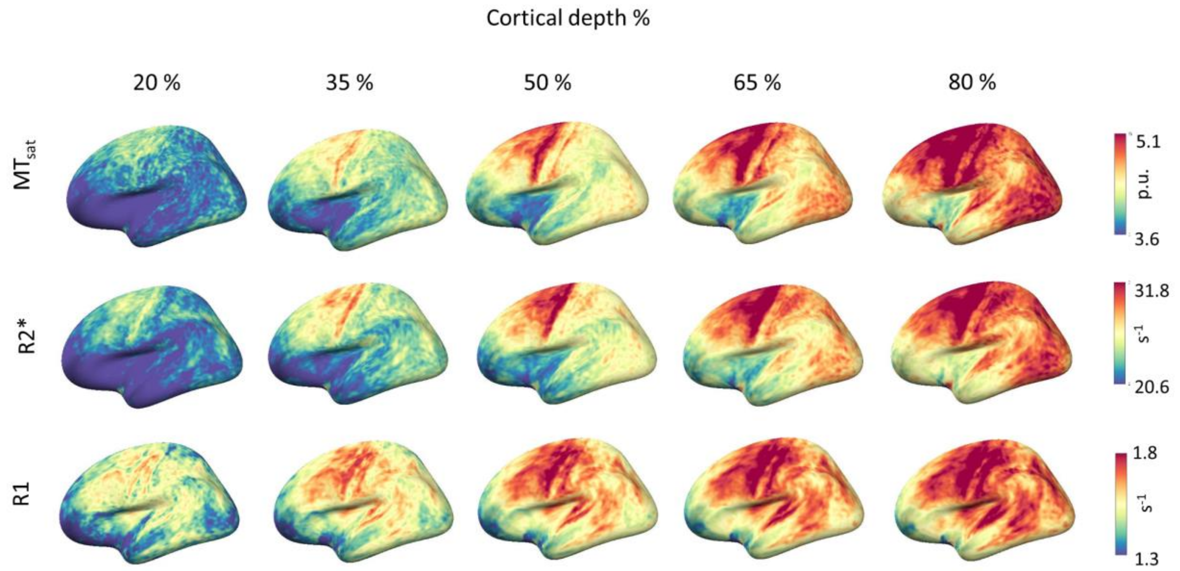

**Supplementary Figure 2:** The group-averaged quantitative parameter maps (R2*,  MTsat, R1) sampled at various cortical depths (in % from pial surface) projected onto the inflated surface of the human *fsaverage* template. Data from 9 adult chimpanzees (age > 17 years) contributed to this plot. Increases in myelination from the pial to the white matter surface are clearly visible across the entire cortex, with the highly myelinated primary areas showing stronger increases.

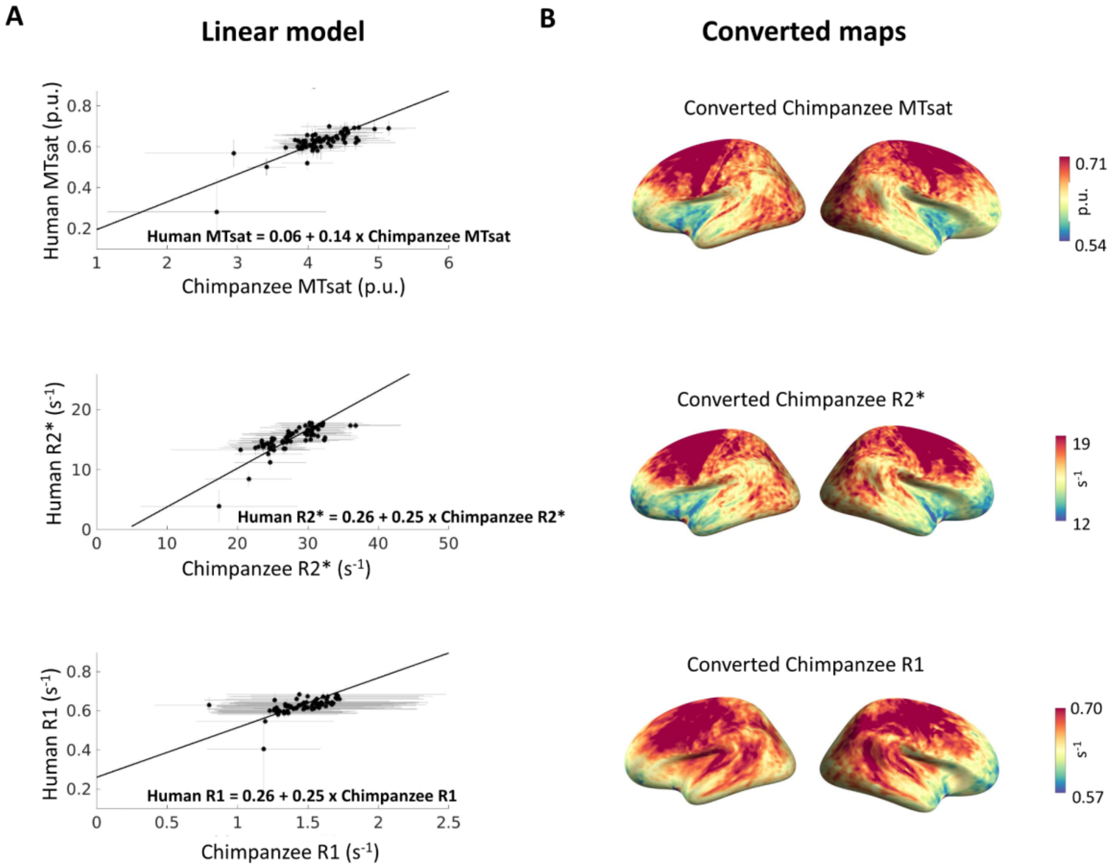

**Supplementary Figure 3**: **A:** For each quantitative parameter, a linear model was determined to account for systematic differences between the *post mortem* chimpanzee data and human *in vivo* data. Mean and standard deviation species values across the 76 BB38 atlas regions contributed to the fitting. Only adult chimpanzee brains contributed to these analyses. The resulting linear model is shown in each plot. **B:** The linear models were applied to harmonize the chimpanzee maps to the range of values found in humans. The resulting group averaged surface projections are shown. Note that the range of values now corresponds to the range of values originally found in humans.

**Supplementary Table 2:** Statistical results of the whole-brain comparison between humans and chimpanzees. Reported are for each analysis (metric and hemisphere), the vertex with the strongest effect size, the size of the cluster and the annotated region of the largest effect. **BB38 atlas[8]:** FA: Primary motor, FB: Premotor cortex, FBA: Ventral precentral gyrus, FC: Premotor cortex, FCBm: Posterior inferior frontal gyrus¸ FDdelta: Prefrontal cortex, FDL: Cingulate cortex¸ FDm: Prefrontal cortex, FDp: Prefrontal cortex, FE: Prefrontal cortex¸ FF: Orbitofrontal cortex, FL: Subgenual cortex, H: Parahippocampal gyrus, IA: Anterior insula¸ IB: Posterior insula¸ LA: Cingulate cortex, LC1: Cingulate cortex, LC2: Cingulate cortex, OA: Peristriate cortex, OB: Peristriate cortex, OC: Primary visual cortex¸ PB: Primary somatosensory¸ PC: Somatosensory cortex¸ PEm: Posterior parietal cortex¸ PF: Posterior parietal cortex, PG: Posterior parietal cortex, PH: Lateral temporal cortex¸ TA: Superior temporal gyrus, TB: Primary auditory, TE1: Middle temporal gyrus¸TE2: Inferior temporal gyrus, TF: Parahippocampal gyrus, TG: Temporal pole

| MTsat left | | | |
|---|---|---|---|
| **Cluster #** | **Vertex max** | **Size ($mm^2$)** | **Annotation** |
| #1 | 19892 | 2029 | FA |
| #2 | 32762 | 685 | FA |
| #3 | 104054 | 234 | TB |
| #4 | 83145 | 152 | unknown |
| #5 | 155610 | 115 | IA |
| #6 | 49910 | 104 | FC |
| MTsat right | | | |
| #1 | 12151 | 640 | FA |
| #2 | 65188 | 550 | FA |
| #3 | 13449 | 152 | FA |
| #4 | 44498 | 114 | unknown |
| R2* left | | | |
| #1 | 140438 | 5025 | FB |
| R2* right | | | |
| #1 | 103565 | 3600 | FB |
| #2 | 50535 | 90 | FC |
| R1 left | | | |

| | | | |
|---|---|---|---|
| #1 | 46045 | 411 | OC |
| No results for R1 right | | | |
| Thickness left | | | |
| #1 | 7560 | 702 | FA |
| #2 | 163764 | 517 | H |
| #3 | 134377 | 484 | LA |
| #4 | 28364 | 440 | TA |
| #5 | 127216 | 439 | OC |
| #6 | 87187 | 377 | OB |
| #7 | 132206 | 310 | FG |
| #8 | 112539 | 275 | OC |
| #9 | 47241 | 202 | PB |
| #10 | 85303 | 194 | IA |
| #11 | 48528 | 116 | PB |
| #12 | 127795 | 115 | OC |
| #13 | 86495 | 110 | PH |
| #14 | 141524 | 105 | IB |
| #15 | 105689 | 97 | PB |
| #16 | 143164 | 96 | FF |
| #17 | 79720 | 89 | OB |
| #18 | 90171 | 74 | OA |
| #19 | 43351 | 69 | TA |

| | | | |
|---|---|---|---|
| #20 | 57252 | 69 | PFD |
| #21 | 133929 | 68 | unknown |
| #22 | 59507 | 67 | FC |
| Thickness right | | | |
| #1 | 3864 | 1272 | TG |
| #2 | 68721 | 789 | OA |
| #3 | 17144 | 579 | OB |
| #4 | 97467 | 450 | TF |
| #5 | 154833 | 411 | LA |
| #6 | 44012 | 408 | OA |
| #7 | 155052 | 391 | FG |
| #8 | 30256 | 311 | PH |
| #9 | 148946 | 303 | OC |
| #10 | 126391 | 278 | OA |
| #11 | 146099 | 149 | FC |
| #12 | 43446 | 140 | LC2 |
| #13 | 70283 | 138 | PH |
| #14 | 28123 | 126 | IB |
| #15 | 64379 | 114 | unknown |
| #16 | 65163 | 112 | FA |
| #17 | 81550 | 111 | PEm |
| #18 | 60336 | 107 | IB |

| | | | |
|---|---|---|---|
| #19 | 5421 | 105 | PG |
| #20 | 111803 | 104 | OC |
| #21 | 72387 | 101 | PEm |
| #22 | 80345 | 95 | FB |
| #23 | 116303 | 75 | LA |
| #24 | 115012 | 68 | TE1 |
| #25 | 53933 | 63 | FB |

**Supplementary Table 3. Modeling results for particularly relevant cortical regions.** Reported are the number of data points that contributed to the modeling (**n**), the model fit as fraction of variance explained by the model (**$R^2$**), the ***p*** value, the constant parameter, the plateau parameter, the % of maximum reached at birth (calculated as 100 * constant / (constant + amplitude)), the rate parameter and the bounds of the 95% confidence interval of the rate parameter. The parameters and regions where the model significantly ($p < .05$) predicted the data are highlighted in gray.

| **R2* (median value across all cortical depths)** | | | | | | | | | | |
|---|---|---|---|---|---|---|---|---|---|---|
| | **n** | **$R^2$** | ***p*** | **Constant** | **Amplitude** | **% at birth** | **Rate** | **Lower CI bound** | **Upper CI bound** | |
| Visual | 17 | 0.82 | < .001 | 13.65 | 25.46 | 35 | 0.03 | -0.02 | 0.09 | |
| Auditory | 17 | 0.8 | < .001 | 13.27 | 26.51 | 31 | 0.03 | -0.03 | 0.08 | |
| Motor | 17 | 0.83 | < .001 | 12.96 | 31.34 | 29 | 0.04 | -0.02 | 0.09 | |
| Somatosensory | 17 | 0.81 | < .001 | 13.32 | 30.09 | 31 | 0.03 | -0.03 | 0.08 | |
| Broca homologue | 17 | 0.82 | < .001 | 12.64 | 25.21 | 33 | 0.02 | -0.03 | 0.08 | |
| Prefrontal | 17 | 0.77 | < .001 | 12.45 | 18.08 | 41 | 0.03 | -0.03 | 0.1 | |
| **MTsat (median value across all cortical depths)** | | | | | | | | | | |
| | **n** | **$R^2$** | ***p*** | **Constant** | **Amplitude** | **% at birth** | **Rate** | **Lower CI bound** | **Upper CI bound** | **Miller model $R^2$** |
| Visual | 17 | 0.74 | < .001 | 2.27 | 2.23 | 50 | 0.48 | 0.09 | 0.87 | 0.65 |
| Auditory | 17 | 0.67 | < .001 | 2.38 | 1.78 | 57 | 0.46 | 0.01 | 0.92 | 0.58 |
| Motor | 17 | 0.86 | < .001 | 2.28 | 2.71 | 46 | 0.38 | 0.14 | 0.61 | 0.81 |
| Somatosensory | 17 | 0.79 | < .001 | 2.34 | 2.19 | 52 | 0.41 | 0.1 | 0.72 | 0.72 |
| Broca homologue | 17 | 0.63 | < .001 | 2.41 | 1.55 | 61 | 0.5 | -0.01 | 1.01 | 0.53 |
| Prefrontal | 17 | 0.53 | 0.001 | 2.39 | 1.56 | 61 | 0.53 | -0.12 | 1.19 | 0.43 |
| **R1 (median value across all cortical depths)** | | | | | | | | | | |
| | **n** | **$R^2$** | ***p*** | **Constant** | **Amplitude** | **% at birth** | **Rate** | **Lower CI bound** | **Upper CI bound** | |
| Visual | 14 | 0.2 | 0.104 | 0.85 | 0.71 | 54 | 0.46 | -0.95 | 1.86 | |
| Auditory | 14 | 0.21 | 0.095 | 0.82 | 0.94 | 47 | 0.32 | -0.8 | 1.44 | |
| Motor | 14 | 0.29 | 0.047 | 0.8 | 0.99 | 45 | 0.34 | -0.61 | 1.29 | |

| | | | | | | | | | |
|---|---|---|---|---|---|---|---|---|---|
| Somatosensory | 14 | 0.26 | 0.06 | 0.81 | 0.93 | 47 | 0.36 | -0.69 | 1.42 |
| Broca homologue | 14 | 0.21 | 0.1 | 0.85 | 0.7 | 55 | 0.35 | -0.85 | 1.55 |
| Prefrontal | 14 | 0.17 | 0.139 | 0.88 | 0.48 | 65 | 0.43 | -1.09 | 1.96 |
| **Thickness** | | | | | | | | | |
| | **n** | **$R^2$** | ***p*** | **Constant** | **Amplitude** | **% at birth** | **Rate** | **Lower CI bound** | **Upper CI bound** |
| Visual | 15 | 0.39 | 0.013 | 2.01 | 5 | | 1.73 | -4.09 | 7.55 |
| Auditory | 15 | 0.19 | 0.105 | 0 | 2.49 | | 0 | -0.23 | 0.24 |
| Motor | 15 | 0.09 | 0.283 | 2.53 | 0.22 | | 0.04 | -0.37 | 0.46 |
| Somatosensory | 15 | 0.29 | 0.036 | 2.19 | 5 | | 1.82 | -5.84 | 9.49 |
| Broca homologue | 15 | 0.15 | 0.149 | 0.09 | 2.83 | | 0 | -0.27 | 0.27 |
| Prefrontal | 15 | 0.53 | 0.002 | 2.92 | 0.86 | | 0.08 | -0.07 | 0.24 |
| **Skewness R2*** | | | | | | | | | |
| | **n** | **$R^2$** | ***p*** | **Constant** | **Amplitude** | **% at birth** | **Slope** | **Lower CI bound** | **Upper CI bound** |
| Visual | 15 | 0 | 0.977 | -0.52 | | | 0 | -0.0022 | 0.0023 |
| Auditory | 15 | 0.3 | 0.035 | -0.49 | | | -0.0038 | -0.0073 | -0.0003 |
| Motor | 15 | 0.23 | 0.072 | -0.55 | | | 0.0031 | -0.0003 | 0.0065 |
| Somatosensory | 15 | 0 | 0.869 | -0.62 | | | 0.0004 | -0.0044 | 0.0051 |
| Broca homologue | 15 | 0.37 | 0.016 | -0.49 | | | -0.0032 | -0.0057 | -0.0007 |
| Prefrontal | 15 | 0.23 | 0.073 | -0.57 | | | -0.0045 | -0.0095 | 0.0005 |
| **Skewness MTsat** | | | | | | | | | |
| | **n** | **$R^2$** | ***p*** | **Constant** | **Amplitude** | **% at birth** | **Slope** | **Lower CI bound** | **Upper CI bound** |
| Visual | 15 | 0.48 | 0.004 | -0.23 | | | -0.0025 | -0.004 | -0.0009 |
| Auditory | 15 | 0.75 | < .001 | -0.21 | | | -0.0043 | -0.0057 | -0.0028 |
| Motor | 15 | 0.36 | 0.017 | -0.25 | | | -0.0023 | -0.0041 | -0.0005 |

| | | | | | | | | | |
|---|---|---|---|---|---|---|---|---|---|
| Somatosensory | 15 | 0.28 | 0.044 | -0.26 | | | -0.0025 | -0.0048 | -0.0001 |
| Broca homologue | 15 | 0.75 | < .001 | -0.18 | | | -0.003 | -0.0041 | -0.002 |
| Prefrontal | 15 | 0.51 | 0.003 | -0.18 | | | -0.0027 | -0.0042 | -0.0011 |

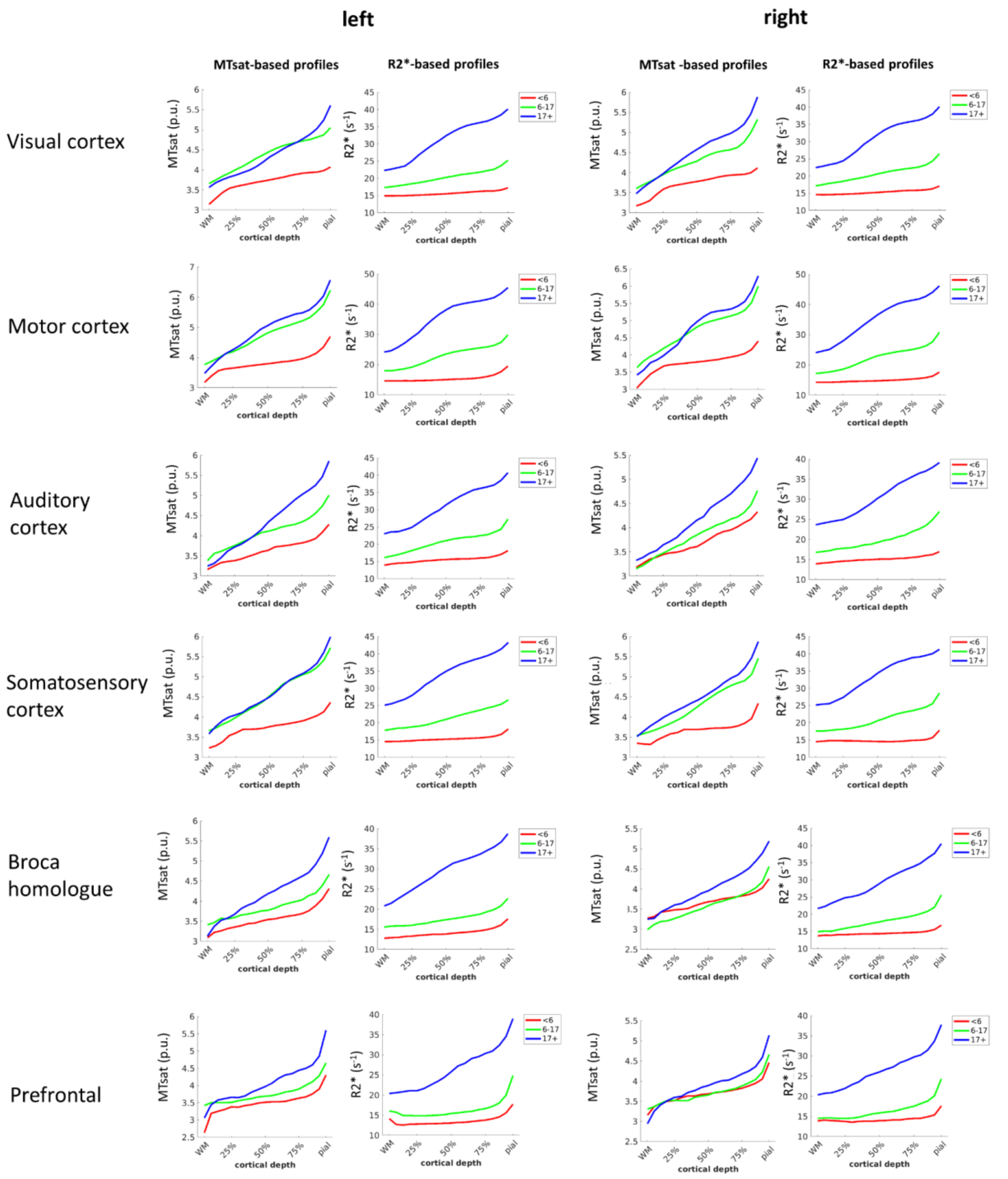

**Supplementary Figure 4**: Age grouped profiles, for young (age < 6 years, N = 3), preadult (age < 17 years, N = 3) and adult (N = 9). Note that profiles are highly reproducible between the left and right hemisphere.

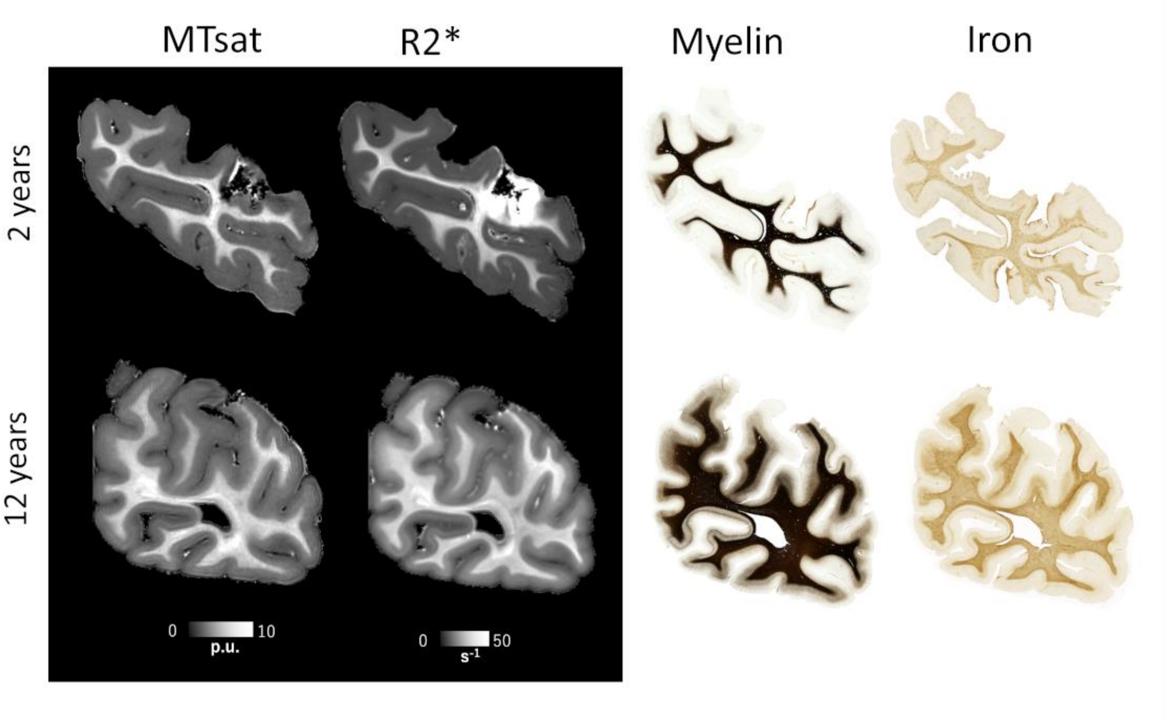

**Supplementary Figure 5**: Histological comparison between the occipital cortex of a 2 year old male chimpanzee who died in the wild and a 12 year old male chimpanzee who died in a sanctuary. Both quantitative MRI and histology indicate more advanced myelination and iron accumulation in the cortex of the 12 year old, and show inter-regional variation. Myelin was visualized with a Gallyas silver stain (note the stain saturated in the white matter). A modified Perls's stain was used to visualize iron.

**Supplementary Table 4. Modeling results for all BB38 cortical regions.** Reported are the number of data points that contributed to the modeling (**n**), the model fit as fraction of variance explained by the model (**$R^2$**), the ***p*** value, the constant parameter, the plateau parameter, the % of maximum reached at birth (calculated as 100 * constant / (constant + amplitude)), the rate parameter and the bounds of the 95% confidence interval of the rate parameter. The parameters and regions where the model significantly ($p < .05$, not corrected for multiple comparisons) predicted the data are highlighted in gray. FA: Primary motor, FB: Premotor cortex, FBA: Ventral precentral gyrus, FC: Premotor cortex, FCBm: Posterior inferior frontal gyrus¸ FDdelta: Prefrontal cortex, FDL: Cingulate cortex¸ FDm: Prefrontal cortex, FDp: Prefrontal cortex, FE: Prefrontal cortex¸ FF: Orbitofrontal cortex, FL: Subgenual cortex, H: Parahippocampal gyrus, IA: Anterior insula¸ IB: Posterior insula¸ LA: Cingulate cortex, LC1: Cingulate cortex, LC2: Cingulate cortex, OA: Peristriate cortex, OB: Peristriate cortex, OC: Primary visual cortex¸ PB: Primary somatosensory¸ PC: Somatosensory cortex¸ PEm: Posterior parietal cortex¸ PF: Posterior parietal cortex, PG: Posterior parietal cortex, PH: Lateral temporal cortex¸ TA: Superior temporal gyrus, TB: Primary auditory, TE1: Middle temporal gyrus¸TE2: Inferior temporal gyrus, TF: Parahippocampal gyrus, TG: Temporal pole

MTsat

| **Region** | **n** | **$R^2$** | ***p*** | **constant** | **amplitude** | **plateau** | **perc** | **time const** | **rate** | **lower rate CI** | **higher rate CI** | **Miller $R^2$** |
|---|---|---|---|---|---|---|---|---|---|---|---|---|
| FA | 17 | 0.86 | <0.001 | 2.28 | 2.71 | 4.98 | 46 | 2.66 | 0.3754 | 0.14 | 0.61 | 0.81 |
| FB | 17 | 0.7 | <0.001 | 2.42 | 2.22 | 4.64 | 52 | 2.9 | 0.3452 | -0.01 | 0.7 | 0.65 |
| FBA | 17 | 0.72 | <0.001 | 2.38 | 2.08 | 4.46 | 53 | 2.4 | 0.4168 | 0.04 | 0.79 | 0.65 |
| FC | 17 | 0.61 | <0.001 | 2.43 | 1.85 | 4.28 | 57 | 2.43 | 0.4114 | -0.07 | 0.89 | 0.56 |
| FCBm | 17 | 0.63 | <0.001 | 2.41 | 1.55 | 3.96 | 61 | 2.02 | 0.4962 | -0.01 | 1.01 | 0.53 |
| FDdelta | 17 | 0.52 | 0.001 | 2.44 | 1.56 | 4 | 61 | 1.99 | 0.5022 | -0.15 | 1.15 | 0.45 |

| | | | | | | | | | | | | |
|---|---|---|---|---|---|---|---|---|---|---|---|---|
| FDL | 17 | 0.52 | 0.001 | 2.38 | 1.41 | 3.8 | 63 | 1.72 | 0.5828 | -0.11 | 1.28 | 0.39 |
| FDm | 17 | 0.53 | 0.001 | 2.39 | 1.56 | 3.95 | 61 | 1.88 | 0.5321 | -0.12 | 1.19 | 0.43 |
| FDp | 17 | 0.59 | <0.001 | 2.4 | 1.54 | 3.94 | 61 | 1.89 | 0.5293 | -0.04 | 1.1 | 0.5 |
| FE | 17 | 0.58 | <0.001 | 2.37 | 1.5 | 3.87 | 61 | 1.78 | 0.5615 | -0.04 | 1.16 | 0.45 |
| FF | 17 | 0.59 | <0.001 | 2.39 | 1.49 | 3.88 | 62 | 1.68 | 0.5962 | -0.02 | 1.21 | 0.45 |
| FG | 17 | 0.52 | 0.001 | 2.35 | 1.6 | 3.95 | 59 | 1.56 | 0.6399 | -0.1 | 1.38 | 0.37 |
| FH | 17 | 0.52 | 0.001 | 2.38 | 1.38 | 3.76 | 63 | 1.64 | 0.6098 | -0.11 | 1.33 | 0.37 |
| FL | 17 | 0.53 | 0.001 | 2.38 | 1.46 | 3.84 | 62 | 1.66 | 0.604 | -0.09 | 1.3 | 0.41 |
| H | 17 | 0.56 | 0.001 | 2.42 | 1.84 | 4.26 | 57 | 3.1 | 0.3223 | -0.14 | 0.79 | 0.53 |
| IA | 17 | 0.54 | 0.001 | 2.42 | 1.27 | 3.69 | 66 | 1.78 | 0.5631 | -0.1 | 1.22 | 0.42 |
| IB | 17 | 0.61 | <0.001 | 2.42 | 1.49 | 3.91 | 62 | 1.84 | 0.5438 | -0.01 | 1.1 | 0.5 |
| LA | 17 | 0.49 | 0.002 | 2.39 | 1.42 | 3.8 | 63 | 1.61 | 0.6227 | -0.15 | 1.39 | 0.35 |
| LC1 | 17 | 0.71 | <0.001 | 2.38 | 1.71 | 4.09 | 58 | 2.17 | 0.4599 | 0.05 | 0.87 | 0.62 |
| LC2 | 17 | 0.64 | <0.001 | 2.41 | 1.67 | 4.08 | 59 | 2.46 | 0.4059 | -0.04 | 0.85 | 0.58 |

| | | | | | | | | | | | | |
|---|---|---|---|---|---|---|---|---|---|---|---|---|
| LE | 17 | 0.62 | <0.001 | 2.98 | 1.67 | 4.65 | 64 | 12.26 | 0.0816 | -0.04 | 0.21 | 0.57 |
| OA | 17 | 0.73 | <0.001 | 2.35 | 2.02 | 4.37 | 54 | 2.22 | 0.4507 | 0.07 | 0.84 | 0.65 |
| OB | 17 | 0.74 | <0.001 | 2.31 | 2.15 | 4.45 | 52 | 2.15 | 0.4643 | 0.09 | 0.84 | 0.67 |
| OC | 17 | 0.74 | <0.001 | 2.27 | 2.23 | 4.5 | 50 | 2.08 | 0.48 | 0.09 | 0.87 | 0.65 |
| PB | 17 | 0.79 | <0.001 | 2.35 | 2.22 | 4.56 | 51 | 2.45 | 0.4087 | 0.1 | 0.71 | 0.74 |
| PC | 17 | 0.79 | <0.001 | 2.34 | 2.19 | 4.54 | 52 | 2.46 | 0.4058 | 0.1 | 0.72 | 0.72 |
| PEm | 17 | 0.81 | <0.001 | 2.33 | 2.16 | 4.5 | 52 | 2.43 | 0.411 | 0.12 | 0.7 | 0.75 |
| PEp | 17 | 0.75 | <0.001 | 2.4 | 2.05 | 4.45 | 54 | 2.67 | 0.375 | 0.04 | 0.71 | 0.69 |
| PF | 17 | 0.74 | <0.001 | 2.37 | 1.95 | 4.33 | 55 | 2.32 | 0.4316 | 0.07 | 0.8 | 0.67 |
| PFD | 17 | 0.71 | <0.001 | 2.37 | 1.85 | 4.22 | 56 | 2.12 | 0.4712 | 0.06 | 0.89 | 0.62 |
| PG | 17 | 0.72 | <0.001 | 2.4 | 1.87 | 4.27 | 56 | 2.36 | 0.4242 | 0.05 | 0.8 | 0.65 |
| PH | 17 | 0.73 | <0.001 | 2.32 | 1.98 | 4.31 | 54 | 2.1 | 0.4769 | 0.08 | 0.88 | 0.65 |
| TA | 17 | 0.66 | <0.001 | 2.37 | 1.72 | 4.09 | 58 | 2.03 | 0.4936 | 0.02 | 0.96 | 0.57 |
| TB | 17 | 0.67 | <0.001 | 2.38 | 1.78 | 4.16 | 57 | 2.15 | 0.4647 | 0.01 | 0.92 | 0.58 |

| | | | | | | | | | | | | |
|---|---|---|---|---|---|---|---|---|---|---|---|---|
| TE1 | 17 | 0.65 | <0.001 | 2.39 | 1.67 | 4.06 | 59 | 2.07 | 0.4824 | -0 | 0.97 | 0.56 |
| TE2 | 17 | 0.66 | <0.001 | 2.33 | 1.8 | 4.13 | 56 | 1.89 | 0.5281 | 0.03 | 1.03 | 0.54 |
| TF | 17 | 0.68 | <0.001 | 2.34 | 1.81 | 4.14 | 56 | 1.86 | 0.5384 | 0.06 | 1.02 | 0.56 |
| TG | 17 | 0.56 | 0.001 | 2.35 | 1.73 | 4.07 | 58 | 1.77 | 0.564 | -0.06 | 1.19 | 0.45 |

R2*

| **Region** | **n** | **$R^2$** | ***p*** | **constant** | **amplitude** | **plateau** | **perc** | **time const** | **rate** | **lower rate CI** | **higher rate CI** | **Miller $R^2$** |
|---|---|---|---|---|---|---|---|---|---|---|---|---|
| FA | 17 | 0.83 | <0.001 | 12.95 | 31.33 | 44.29 | 29 | 25.43 | 0.0393 | -0.02 | 0.09 | 0.76 |
| FB | 17 | 0.87 | <0.001 | 12.21 | 27.59 | 39.8 | 31 | 25.16 | 0.0397 | -0.01 | 0.09 | 0.8 |
| FBA | 17 | 0.84 | <0.001 | 12.72 | 29.32 | 42.04 | 30 | 32.87 | 0.0304 | -0.02 | 0.08 | 0.76 |
| FC | 17 | 0.86 | <0.001 | 12.4 | 25.49 | 37.89 | 33 | 30.06 | 0.0333 | -0.01 | 0.08 | 0.79 |
| FCBm | 17 | 0.82 | <0.001 | 12.64 | 25.2 | 37.84 | 33 | 41.41 | 0.0241 | -0.03 | 0.08 | 0.75 |
| FDdelta | 17 | 0.82 | <0.001 | 12.25 | 19.07 | 31.32 | 39 | 25.09 | 0.0399 | -0.02 | 0.1 | 0.78 |

| | | | | | | | | | | | | |
|---|---|---|---|---|---|---|---|---|---|---|---|---|
| FDL | 17 | 0.73 | <0.001 | 12.93 | 23.27 | 36.21 | 36 | 54.77 | 0.0183 | -0.05 | 0.08 | 0.67 |
| FDm | 17 | 0.77 | <0.001 | 12.45 | 18.07 | 30.53 | 41 | 29.92 | 0.0334 | -0.03 | 0.1 | 0.71 |
| FDp | 17 | 0.76 | <0.001 | 12.73 | 21.32 | 34.05 | 37 | 41.2 | 0.0243 | -0.04 | 0.09 | 0.7 |
| FE | 17 | 0.69 | <0.001 | 12.77 | 19.23 | 31.99 | 40 | 42.2 | 0.0237 | -0.05 | 0.1 | 0.63 |
| FF | 17 | 0.73 | <0.001 | 12.89 | 18.95 | 31.85 | 40 | 33.76 | 0.0296 | -0.04 | 0.1 | 0.69 |
| FG | 17 | 0.66 | <0.001 | 13.04 | 13.94 | 26.99 | 48 | 23.4 | 0.0427 | -0.04 | 0.13 | 0.62 |
| FH | 17 | 0.63 | <0.001 | 13.13 | 20.78 | 33.91 | 39 | 53.54 | 0.0187 | -0.07 | 0.1 | 0.58 |
| FL | 17 | 0.73 | <0.001 | 13.4 | 103.74 | 117.14 | 11 | 305.85 | 0.0033 | -0.06 | 0.07 | 0.63 |
| H | 17 | 0.74 | <0.001 | 11.55 | 19.11 | 30.67 | 38 | 18.64 | 0.0536 | -0.02 | 0.13 | 0.75 |
| IA | 17 | 0.71 | <0.001 | 13.08 | 19.49 | 32.57 | 40 | 35.57 | 0.0281 | -0.05 | 0.1 | 0.66 |
| IB | 17 | 0.74 | <0.001 | 13.8 | 274.08 | 287.88 | 5 | 734.4 | 0.0014 | -0.06 | 0.06 | 0.63 |
| LA | 17 | 0.73 | <0.001 | 13.33 | 38.97 | 52.3 | 25 | 101.82 | 0.0098 | -0.05 | 0.07 | 0.66 |
| LC1 | 17 | 0.78 | <0.001 | 13.31 | 28.5 | 41.81 | 32 | 50.73 | 0.0197 | -0.04 | 0.08 | 0.7 |
| LC2 | 17 | 0.78 | <0.001 | 13.1 | 31.5 | 44.6 | 29 | 56.36 | 0.0177 | -0.04 | 0.08 | 0.7 |

| | | | | | | | | | | | | |
|---|---|---|---|---|---|---|---|---|---|---|---|---|
| LE | 17 | 0.82 | <0.001 | 12.92 | 26.35 | 39.27 | 33 | 26.82 | 0.0373 | -0.02 | 0.09 | 0.76 |
| OA | 17 | 0.85 | <0.001 | 13.03 | 24.22 | 37.25 | 35 | 27.42 | 0.0365 | -0.01 | 0.09 | 0.78 |
| OB | 17 | 0.83 | <0.001 | 13.26 | 24.47 | 37.74 | 35 | 28.03 | 0.0357 | -0.02 | 0.09 | 0.76 |
| OC | 17 | 0.82 | <0.001 | 13.65 | 25.48 | 39.12 | 35 | 32.6 | 0.0307 | -0.02 | 0.09 | 0.74 |
| PB | 17 | 0.8 | <0.001 | 13.69 | 32.2 | 45.89 | 30 | 42.31 | 0.0236 | -0.03 | 0.08 | 0.72 |
| PC | 17 | 0.81 | <0.001 | 13.32 | 30.08 | 43.4 | 31 | 36.13 | 0.0277 | -0.03 | 0.08 | 0.74 |
| PEm | 17 | 0.82 | <0.001 | 12.86 | 25.3 | 38.16 | 34 | 25.85 | 0.0387 | -0.02 | 0.09 | 0.76 |
| PEp | 17 | 0.85 | <0.001 | 12.7 | 27.73 | 40.44 | 31 | 30.5 | 0.0328 | -0.02 | 0.08 | 0.78 |
| PF | 17 | 0.82 | <0.001 | 13.32 | 29.12 | 42.44 | 31 | 38.51 | 0.026 | -0.03 | 0.08 | 0.74 |
| PFD | 17 | 0.83 | <0.001 | 13.2 | 29.11 | 42.31 | 31 | 41.33 | 0.0242 | -0.03 | 0.08 | 0.75 |
| PG | 17 | 0.81 | <0.001 | 13.13 | 26.93 | 40.06 | 33 | 36.02 | 0.0278 | -0.03 | 0.08 | 0.74 |
| PH | 17 | 0.82 | <0.001 | 13.27 | 27.3 | 40.57 | 33 | 38.24 | 0.0262 | -0.03 | 0.08 | 0.74 |
| TA | 17 | 0.81 | <0.001 | 13.19 | 29.06 | 42.25 | 31 | 49.62 | 0.0202 | -0.03 | 0.07 | 0.73 |
| TB | 17 | 0.8 | <0.001 | 13.27 | 26.52 | 39.79 | 33 | 37.33 | 0.0268 | -0.03 | 0.08 | 0.73 |

| | | | | | | | | | | | | |
|---|---|---|---|---|---|---|---|---|---|---|---|---|
| TE1 | 17 | 0.79 | <0.001 | 13.02 | 25.02 | 38.04 | 34 | 44.33 | 0.0226 | -0.03 | 0.08 | 0.73 |
| TE2 | 17 | 0.78 | <0.001 | 13.28 | 30.15 | 43.43 | 31 | 58.6 | 0.0171 | -0.04 | 0.08 | 0.69 |
| TF | 17 | 0.77 | <0.001 | 13.45 | 26.73 | 40.18 | 33 | 45.73 | 0.0219 | -0.04 | 0.08 | 0.7 |
| TG | 17 | 0.75 | <0.001 | 13.68 | 34.77 | 48.45 | 28 | 91.64 | 0.0109 | -0.05 | 0.07 | 0.66 |

R1

| **Region** | **n** | **$R^2$** | ***p*** | **constant** | **amplitude** | **plateau** | **perc** | **time const** | **rate** | **lower rate CI** | **higher rate CI** | **Miller $R^2$** |
|---|---|---|---|---|---|---|---|---|---|---|---|---|
| FA | 14 | 0.29 | 0.047 | 0.8 | 0.99 | 1.79 | 45 | 2.94 | 0.3403 | -0.61 | 1.29 | 0.43 |
| FB | 14 | 0.28 | 0.053 | 0.84 | 0.79 | 1.63 | 51 | 2.93 | 0.3413 | -0.64 | 1.33 | 0.39 |
| FBA | 14 | 0.25 | 0.069 | 0.84 | 0.87 | 1.71 | 49 | 2.89 | 0.3457 | -0.72 | 1.41 | 0.37 |
| FC | 14 | 0.26 | 0.06 | 0.85 | 0.73 | 1.58 | 54 | 2.95 | 0.3384 | -0.67 | 1.35 | 0.38 |
| FCBm | 14 | 0.21 | 0.1 | 0.85 | 0.7 | 1.55 | 55 | 2.85 | 0.3508 | -0.85 | 1.56 | 0.33 |
| FDdelta | 14 | 0.19 | 0.114 | 0.88 | 0.6 | 1.49 | 59 | 2.66 | 0.3759 | -0.93 | 1.69 | 0.3 |

| | | | | | | | | | | | | |
|---|---|---|---|---|---|---|---|---|---|---|---|---|
| FDL | 14 | 0.15 | 0.178 | 0.89 | 0.45 | 1.34 | 66 | 2.82 | 0.3545 | -1.15 | 1.86 | 0.25 |
| FDm | 14 | 0.17 | 0.139 | 0.88 | 0.48 | 1.36 | 65 | 2.31 | 0.4333 | -1.09 | 1.96 | 0.32 |
| FDp | 14 | 0.18 | 0.135 | 0.88 | 0.56 | 1.43 | 61 | 2.47 | 0.4051 | -1.05 | 1.86 | 0.31 |
| FE | 14 | 0.15 | 0.175 | 0.88 | 0.47 | 1.35 | 65 | 2.21 | 0.4519 | -1.26 | 2.17 | 0.36 |
| FF | 14 | 0.17 | 0.146 | 0.87 | 0.64 | 1.51 | 58 | 2.53 | 0.3949 | -1.08 | 1.87 | 0.34 |
| FG | 14 | 0.13 | 0.199 | 0.85 | 0.58 | 1.43 | 60 | 2.16 | 0.4639 | -1.38 | 2.31 | 0.42 |
| FH | 14 | 0.14 | 0.189 | 0.87 | 0.48 | 1.35 | 64 | 2.55 | 0.3917 | -1.25 | 2.03 | 0.31 |
| FL | 14 | 0.15 | 0.165 | 0.87 | 0.6 | 1.47 | 59 | 2.3 | 0.4345 | -1.2 | 2.07 | 0.33 |
| H | 14 | 0.18 | 0.129 | 0.84 | 0.56 | 1.4 | 60 | 2.73 | 0.3658 | -0.98 | 1.71 | 0.38 |
| IA | 14 | 0.18 | 0.131 | 0.84 | 0.76 | 1.6 | 53 | 3.15 | 0.318 | -0.93 | 1.57 | 0.29 |
| IB | 14 | 0.22 | 0.087 | 0.84 | 0.89 | 1.73 | 48 | 3 | 0.3338 | -0.79 | 1.45 | 0.37 |
| LA | 14 | 0.17 | 0.146 | 0.87 | 0.56 | 1.43 | 61 | 2.91 | 0.3435 | -1.02 | 1.71 | 0.3 |
| LC1 | 14 | 0.2 | 0.105 | 0.84 | 0.73 | 1.57 | 53 | 2.96 | 0.3377 | -0.86 | 1.54 | 0.35 |
| LC2 | 14 | 0.16 | 0.156 | 0.86 | 0.56 | 1.42 | 61 | 3.34 | 0.2992 | -0.99 | 1.59 | 0.28 |

| | | | | | | | | | | | | |
|---|---|---|---|---|---|---|---|---|---|---|---|---|
| LE | 14 | 0.18 | 0.132 | 0.83 | 0.68 | 1.51 | 55 | 3.35 | 0.2986 | -0.91 | 1.51 | 0.29 |
| OA | 14 | 0.22 | 0.091 | 0.84 | 0.76 | 1.6 | 53 | 2.59 | 0.3867 | -0.85 | 1.62 | 0.39 |
| OB | 14 | 0.21 | 0.101 | 0.85 | 0.71 | 1.56 | 54 | 2.44 | 0.4093 | -0.91 | 1.72 | 0.39 |
| OC | 14 | 0.2 | 0.104 | 0.85 | 0.71 | 1.56 | 54 | 2.2 | 0.454 | -0.95 | 1.86 | 0.4 |
| PB | 14 | 0.26 | 0.063 | 0.8 | 1 | 1.8 | 45 | 3.03 | 0.3305 | -0.68 | 1.34 | 0.4 |
| PC | 14 | 0.26 | 0.06 | 0.81 | 0.94 | 1.75 | 46 | 2.74 | 0.3647 | -0.69 | 1.42 | 0.41 |
| PEm | 14 | 0.24 | 0.075 | 0.82 | 0.86 | 1.67 | 49 | 2.81 | 0.3564 | -0.75 | 1.47 | 0.39 |
| PEp | 14 | 0.25 | 0.068 | 0.83 | 0.83 | 1.66 | 50 | 2.91 | 0.3431 | -0.71 | 1.4 | 0.38 |
| PF | 14 | 0.25 | 0.069 | 0.82 | 0.94 | 1.76 | 47 | 2.91 | 0.344 | -0.72 | 1.41 | 0.38 |
| PFD | 14 | 0.23 | 0.081 | 0.83 | 0.86 | 1.7 | 49 | 2.8 | 0.3572 | -0.78 | 1.49 | 0.38 |
| PG | 14 | 0.24 | 0.073 | 0.83 | 0.89 | 1.72 | 48 | 2.86 | 0.3491 | -0.74 | 1.44 | 0.37 |
| PH | 14 | 0.22 | 0.091 | 0.83 | 0.82 | 1.66 | 50 | 2.6 | 0.384 | -0.85 | 1.61 | 0.37 |
| TA | 14 | 0.2 | 0.106 | 0.84 | 0.82 | 1.66 | 51 | 2.85 | 0.3512 | -0.88 | 1.58 | 0.32 |
| TB | 14 | 0.21 | 0.095 | 0.82 | 0.94 | 1.76 | 47 | 3.14 | 0.3182 | -0.8 | 1.44 | 0.32 |

| | | | | | | | | | | | | |
|---|---|---|---|---|---|---|---|---|---|---|---|---|
| TE1 | 14 | 0.2 | 0.109 | 0.83 | 0.77 | 1.6 | 52 | 2.9 | 0.3454 | -0.88 | 1.57 | 0.34 |
| TE2 | 14 | 0.18 | 0.125 | 0.86 | 0.66 | 1.52 | 57 | 2.22 | 0.4506 | -1.05 | 1.95 | 0.4 |
| TF | 14 | 0.16 | 0.163 | 0.87 | 0.6 | 1.47 | 59 | 2.57 | 0.3898 | -1.14 | 1.92 | 0.35 |
| TG | 14 | 0.17 | 0.142 | 0.85 | 0.62 | 1.47 | 58 | 2.43 | 0.4123 | -1.08 | 1.91 | 0.38 |

Thickness

| **Region** | **n** | **$R^2$** | ***p*** | **constant** | **amplitude** | **plateau** | **time const** | **rate** | **lower rate CI** | **higher rate CI** | **Miller $R^2$** |
|---|---|---|---|---|---|---|---|---|---|---|---|
| FA | 15 | 0.09 | 0.283 | 2.53 | 0.22 | 2.75 | 22.87 | 0.0437 | -0.37 | 0.46 | 0.15 |
| FB | 15 | 0.36 | 0.018 | 2.84 | 0.42 | 3.26 | 15.03 | 0.0665 | -0.13 | 0.26 | 0.34 |
| FBA | 15 | 0.19 | 0.107 | 2.36 | 0.51 | 2.87 | 26.39 | 0.0379 | -0.22 | 0.3 | 0.26 |
| FC | 15 | 0.32 | 0.029 | 2.6 | 0.37 | 2.98 | 21.46 | 0.0466 | -0.15 | 0.24 | 0.49 |
| FCBm | 15 | 0.15 | 0.149 | 0.09 | 2.83 | 2.92 | 568.79 | 0.0018 | -0.27 | 0.27 | 0.15 |
| FDdelta | 15 | 0.13 | 0.183 | 2.47 | 0.24 | 2.71 | 14.08 | 0.071 | -0.32 | 0.46 | 0.15 |

| | | | | | | | | | | | |
|---|---|---|---|---|---|---|---|---|---|---|---|
| FDL | 15 | 0.42 | 0.009 | 3.02 | 1.14 | 4.17 | 13.04 | 0.0767 | -0.11 | 0.26 | 0.45 |
| FDm | 15 | 0.53 | 0.002 | 2.92 | 0.86 | 3.78 | 11.82 | 0.0846 | -0.07 | 0.24 | 0.57 |
| FDp | 15 | 0.47 | 0.005 | 2.58 | 0.65 | 3.23 | 16.67 | 0.06 | -0.09 | 0.21 | 0.42 |
| FE | 15 | 0.63 | <0.001 | 2.64 | 5 | 7.64 | 0.88 | 1.1368 | -0.94 | 3.21 | 0.39 |
| FF | 15 | 0.24 | 0.066 | 2.78 | 0.37 | 3.15 | 23.63 | 0.0423 | -0.19 | 0.27 | 0.21 |
| FG | 15 | 0.16 | 0.138 | 2.84 | 1.52 | 4.35 | 1.62 | 0.6192 | -2.69 | 3.93 | 0.16 |
| FH | 15 | 0.06 | 0.385 | 3.43 | 0.49 | 3.92 | 9.08 | 0.1102 | -0.7 | 0.92 | 0.05 |
| FL | 15 | 0.02 | 0.626 | 2.85 | 0.32 | 3.17 | 134.58 | 0.0074 | -0.82 | 0.83 | 0 |
| H | 15 | 0.12 | 0.21 | 2.05 | 1.25 | 3.31 | 1.89 | 0.5281 | -2.87 | 3.92 | 0.17 |
| IA | 15 | 0.01 | 0.691 | 2.73 | 0.08 | 2.81 | 22.48 | 0.0445 | -1.1 | 1.19 | 0.01 |
| IB | 15 | 0.3 | 0.035 | 2.09 | 0.44 | 2.53 | 81.23 | 0.0123 | -0.16 | 0.19 | 0.38 |
| LA | 15 | 0.77 | <0.001 | 2.73 | 1.06 | 3.79 | 17.58 | 0.0569 | -0.02 | 0.13 | 0.75 |
| LC1 | 15 | 0.49 | 0.004 | 2.4 | 0.47 | 2.87 | 24.11 | 0.0415 | -0.09 | 0.17 | 0.49 |
| LC2 | 15 | 0.78 | <0.001 | 2.35 | 2.01 | 4.36 | 2.79 | 0.3588 | -0.08 | 0.8 | 0.75 |

| | | | | | | | | | | | |
|---|---|---|---|---|---|---|---|---|---|---|---|
| LE | 15 | 0.68 | <0.001 | 1.71 | 1.76 | 3.48 | 4.72 | 0.212 | -0.08 | 0.51 | 0.67 |
| OA | 15 | 0.28 | 0.044 | 2.24 | 0.32 | 2.55 | 7.02 | 0.1424 | -0.28 | 0.56 | 0.26 |
| OB | 15 | 0.56 | 0.001 | 2.11 | 5 | 7.11 | 0.65 | 1.5397 | -1.84 | 4.92 | 0.32 |
| OC | 15 | 0.39 | 0.013 | 2.01 | 5 | 7.01 | 0.58 | 1.7284 | -4.09 | 7.55 | 0.1 |
| PB | 15 | 0.02 | 0.575 | 3.15 | -1.17 | 1.98 | 1012 | 0.001 | -0.71 | 0.71 | 0.28 |
| PC | 15 | 0.29 | 0.036 | 2.19 | 5 | 7.19 | 0.55 | 1.8237 | -5.84 | 9.49 | 0.06 |
| PEm | 15 | 0.54 | 0.002 | 2.24 | 5 | 7.24 | 0.67 | 1.5006 | -1.99 | 4.99 | 0.24 |
| PEp | 15 | 0.34 | 0.023 | 2.21 | 0.43 | 2.64 | 10.07 | 0.0993 | -0.16 | 0.36 | 0.38 |
| PF | 15 | 0.47 | 0.005 | 2.4 | 5 | 7.4 | 0.62 | 1.6235 | -2.62 | 5.87 | 0.31 |
| PFD | 15 | 0.17 | 0.123 | 2.46 | 0.27 | 2.73 | 28.29 | 0.0353 | -0.24 | 0.31 | 0.13 |
| PG | 15 | 0.21 | 0.088 | 2.38 | 0.22 | 2.6 | 11.78 | 0.0849 | -0.24 | 0.41 | 0.28 |
| PH | 15 | 0.56 | 0.001 | 2.48 | 1.49 | 3.97 | 1.24 | 0.8064 | -0.85 | 2.46 | 0.45 |
| TA | 15 | 0.08 | 0.298 | 1.55 | 0.99 | 2.55 | 421.75 | 0.0024 | -0.38 | 0.38 | 0.08 |
| TB | 15 | 0.19 | 0.105 | 0 | 2.49 | 2.49 | 466 | 0.0021 | -0.23 | 0.24 | 0.23 |

| | | | | | | | | | | | |
|---|---|---|---|---|---|---|---|---|---|---|---|
| TE1 | 15 | 0.1 | 0.257 | 0.97 | 1.67 | 2.63 | 419.69 | 0.0024 | -0.34 | 0.35 | 0.2 |
| TE2 | 15 | 0.08 | 0.308 | 2.73 | 0.38 | 3.11 | 2.34 | 0.4277 | -2.95 | 3.8 | 0.07 |
| TF | 15 | 0.35 | 0.02 | 2.16 | 0.63 | 2.79 | 6.28 | 0.1592 | -0.24 | 0.56 | 0.4 |
| TG | 15 | 0.22 | 0.074 | 5 | -2.3 | 2.7 | 252.76 | 0.004 | -0.21 | 0.22 | 0.34 |

Skewness MTsat

| **Region** | **n** | **$R^2$** | ***p*** | **constant** | **slope** |
|---|---|---|---|---|---|
| FA | 15 | 0.36 | 0.017 | -0.25 | -0.0023 |
| FB | 15 | 0.55 | 0.002 | -0.22 | -0.0026 |
| FBA | 15 | 0.63 | 0 | -0.22 | -0.0029 |
| FC | 15 | 0.75 | 0 | -0.17 | -0.0035 |
| FCBm | 15 | 0.75 | 0 | -0.18 | -0.003 |
| FDdelta | 15 | 0.65 | 0 | -0.16 | -0.0032 |
| FDL | 15 | 0.59 | 0.001 | -0.17 | -0.0021 |

| | | | | | |
|---|---|---|---|---|---|
| FDm | 15 | 0.51 | 0.003 | -0.18 | -0.0027 |
| FDp | 15 | 0.55 | 0.002 | -0.17 | -0.0028 |
| FE | 15 | 0.56 | 0.001 | -0.17 | -0.0025 |
| FF | 15 | 0.6 | 0.001 | -0.15 | -0.003 |
| FG | 15 | 0.17 | 0.124 | -0.17 | -0.0017 |
| FH | 15 | 0.33 | 0.025 | -0.17 | -0.0016 |
| FL | 15 | 0.48 | 0.004 | -0.11 | -0.0024 |
| H | 15 | 0.41 | 0.01 | -0.14 | -0.0024 |
| IA | 15 | 0.63 | 0 | -0.13 | -0.0029 |
| IB | 15 | 0.72 | 0 | -0.17 | -0.0029 |
| LA | 15 | 0.71 | 0 | -0.18 | -0.0028 |
| LC1 | 15 | 0.59 | 0.001 | -0.18 | -0.0031 |
| LC2 | 15 | 0.62 | 0.001 | -0.14 | -0.0039 |
| LE | 15 | 0.13 | 0.189 | -0.12 | -0.0022 |

| | | | | | |
|---|---|---|---|---|---|
| OA | 15 | 0.65 | 0 | -0.21 | -0.0033 |
| OB | 15 | 0.5 | 0.003 | -0.25 | -0.0025 |
| OC | 15 | 0.48 | 0.004 | -0.23 | -0.0025 |
| PB | 15 | 0.38 | 0.014 | -0.23 | -0.0024 |
| PC | 15 | 0.28 | 0.044 | -0.26 | -0.0025 |
| PEm | 15 | 0.36 | 0.017 | -0.23 | -0.0026 |
| PEp | 15 | 0.6 | 0.001 | -0.19 | -0.0035 |
| PF | 15 | 0.6 | 0.001 | -0.18 | -0.0032 |
| PFD | 15 | 0.64 | 0 | -0.2 | -0.0035 |
| PG | 15 | 0.61 | 0.001 | -0.17 | -0.0037 |
| PH | 15 | 0.46 | 0.005 | -0.22 | -0.003 |
| TA | 15 | 0.7 | 0 | -0.18 | -0.0037 |
| TB | 15 | 0.75 | 0 | -0.21 | -0.0043 |
| TE1 | 15 | 0.63 | 0 | -0.16 | -0.0031 |

| | | | | | |
|---|---|---|---|---|---|
| TE2 | 15 | 0.62 | 0 | -0.17 | -0.0025 |
| TF | 15 | 0.6 | 0.001 | -0.18 | -0.0033 |
| TG | 15 | 0.01 | 0.666 | -0.21 | -0.0004 |

Skewness R2*

| **Region** | **n** | **$R^2$** | ***p*** | **constant** | **slope** |
|---|---|---|---|---|---|
| FA | 15 | 0.23 | 0.072 | -0.55 | 0.0031 |
| FB | 15 | 0.14 | 0.167 | -0.6 | 0.0029 |
| FBA | 15 | 0.02 | 0.611 | -0.53 | 0.0009 |
| FC | 15 | 0.07 | 0.337 | -0.49 | -0.0022 |
| FCBm | 15 | 0.37 | 0.016 | -0.49 | -0.0032 |
| FDdelta | 15 | 0.34 | 0.021 | -0.47 | -0.0049 |
| FDL | 15 | 0.46 | 0.006 | -0.5 | -0.0054 |
| FDm | 15 | 0.23 | 0.073 | -0.57 | -0.0045 |

| | | | | | |
|---|---|---|---|---|---|
| FDp | 15 | 0.4 | 0.012 | -0.46 | -0.0052 |
| FE | 15 | 0.51 | 0.003 | -0.5 | -0.0068 |
| FF | 15 | 0.5 | 0.003 | -0.43 | -0.0069 |
| FG | 15 | 0.52 | 0.002 | -0.47 | -0.0072 |
| FH | 15 | 0.14 | 0.164 | -0.5 | -0.0037 |
| FL | 15 | 0.52 | 0.002 | -0.4 | -0.0084 |
| H | 15 | 0.53 | 0.002 | -0.36 | -0.007 |
| IA | 15 | 0.37 | 0.016 | -0.29 | -0.0093 |
| IB | 15 | 0.41 | 0.01 | -0.49 | -0.0046 |
| LA | 15 | 0.54 | 0.002 | -0.56 | -0.0062 |
| LC1 | 15 | 0.35 | 0.021 | -0.46 | -0.0052 |
| LC2 | 15 | 0.13 | 0.196 | -0.52 | -0.0034 |
| LE | 15 | <0.001 | 0.83 | -0.35 | -0.0005 |
| OA | 15 | 0.17 | 0.121 | -0.53 | -0.0023 |

| | | | | | |
|---|---|---|---|---|---|
| OB | 15 | 0.06 | 0.363 | -0.56 | -0.0012 |
| OC | 15 | <0.001 | 0.977 | -0.52 | 0 |
| PB | 15 | 0.12 | 0.208 | -0.45 | -0.0029 |
| PC | 15 | <0.001 | 0.869 | -0.62 | 0.0004 |
| PEm | 15 | 0.07 | 0.359 | -0.51 | -0.0018 |
| PEp | 15 | 0.1 | 0.262 | -0.47 | -0.0026 |
| PF | 15 | 0.16 | 0.139 | -0.48 | -0.0026 |
| PFD | 15 | 0.17 | 0.126 | -0.51 | -0.0021 |
| PG | 15 | 0.23 | 0.07 | -0.44 | -0.0042 |
| PH | 15 | 0.04 | 0.472 | -0.56 | -0.0014 |
| TA | 15 | 0.27 | 0.046 | -0.5 | -0.0041 |
| TB | 15 | 0.3 | 0.035 | -0.49 | -0.0038 |
| TE1 | 15 | 0.35 | 0.021 | -0.45 | -0.0046 |
| TE2 | 15 | 0.36 | 0.019 | -0.45 | -0.0036 |

| | | | | | |
|---|---|---|---|---|---|
| TF | 15 | 0.3 | 0.034 | -0.5 | -0.0043 |
| TG | 15 | 0.65 | 0 | -0.45 | -0.007 |
